# AI-qualizing Science

**DOI:** 10.1101/2025.02.11.637417

**Authors:** Anantha Divakaruni, Francois Bares, Ludovic Phalippou

**Affiliations:** Department of Economics, University of Bergen, Norway; Saïd Business School, University of Oxford, UK

## Abstract

Scientific discovery remains highly concentrated among elite universities due to unequal access to infrastructure, expertise, and collaboration networks. We investigate whether artificial intelligence can mitigate these disparities by studying AlphaFold, a deep learning system awarded the 2024 Nobel Prize in Chemistry for its transformative impact on protein structure predictions. Using publication data from top journals and universities worldwide, we show that lower-ranked institutions increased their share of high-impact protein research by up to five percentage points within two years of AlphaFold’s public release. These gains are specific to protein domains and absent in non-protein fields or lower-tier journals. We further document enhanced research novelty, directional pivoting, and citation impact among lower-tier institutions, alongside reduced dependence on collaborations with top-ranked partners. By broadening participation in frontier protein science, AlphaFold exemplifies how open-access AI tools can disrupt entrenched hierarchies in research. This democratizing effect has far-reaching implications as similar AI systems emerge in other complex domains such as genomics, materials science, and climate modeling.

## Introduction

Although several articles explore the influence of AI on low and medium-skill activities such as image recognition [1], programming [2], writing [3], and customer support [4], there is limited evidence on the impact of AI on productivity within highly specialized sectors. Filling this gap is crucial due to AI rapidly advancing from automating relatively straightforward tasks (e.g., chatbots, code generators) to engaging in complex, high-skill areas that demand substantial expertise and research infrastructure.

The most profound impact of AI in a specialized field has arguably been observed recently within the life sciences. An AI tool called AlphaFold, developed by Google DeepMind, has achieved very high accuracy levels in predicting three-dimensional (3D) protein structures that closely match those of experimental methods [5–7]. As precise protein structures are crucial for understanding protein functions, designing therapeutics, and investigating biological pathways, AlphaFold is widely recognized as a significant advance in structural biology and the broader domain of protein research. Consequently, its creators were honored with the 2023 Breakthrough Prize in Life Sciences and the 2024 Nobel Prize in Chemistry.

In the domain of protein research, where groundbreaking discoveries often depend on access to state-of-the-art resources and specialized knowledge, the *Matthew* effect — the tendency for resources, recognition, and influence to accrue to those who already have them — is particularly evident [8–11]. Before the release of AlphaFold, researchers in the top 10% universities published 55% of all articles in leading journals and secured 50% of the research grants [12, 13]. In addition, top scientists often serve as editors of journals and grant reviewers, which may favor research conducted by scientists affiliated with similarly prestigious institutions [14–16].

A tool such as AlphaFold might be expected to widen existing disparities, enabling elite institutions to exploit its technical advantages more rapidly and thoroughly due to their superior human capital, infrastructure, and entrenched influence within the publication ecosystem. Yet, by sharply lowering the technical and computational demands of protein structure prediction, AlphaFold may instead help level the playing field. When broadly accessible and user-friendly, it can empower researchers at lower-ranked institutions, traditionally constrained in resource-intensive fields, to compete more effectively for space in top-tier journals.

This study investigates whether AlphaFold has modified the existing stratification of high-impact scientific publications across institutions. Universities are classified into quartiles according to their overall publication shares in top-tier journals prior to the release of AlphaFold. By examining the changes in each quartile’s proportion of publications in leading journals, we assess whether the innovative features and open access associated with AlphaFold have facilitated more equitable access to premier scientific venues or have perpetuated existing disparities.

Our analysis shows that following the introduction of AlphaFold, universities in the bottom quartile (ranked 137^th^ to 500^th^ pre-AlphaFold) experienced notable increases in top-tier publications. Similar, though smaller, gains are observed for institutions in the second and third quartiles. In contrast, top-quartile universities saw a decline in their market share of high-impact publications after AlphaFold’s release. These patterns suggest that artificial intelligence can reduce entry barriers in resource-intensive scientific fields and disrupt established hierarchies in knowledge production, contributing to a more balanced distribution of scientific opportunity.

We also present additional evidence suggesting that changes in research practices may have accompanied these shifts in publication patterns. In particular, lower-quartile institutions exhibit greater pivots in research direction, engage more frequently with less commonly used concepts (more novelty), and experience some increases in disruptiveness. While these patterns are most pronounced in structural biology, we note similar trends in non-structural protein domains. Collaboration dynamics also evolve: institutions with stronger structural biology capacity become less reliant on top-tier partners, while those with limited in-house expertise increase such collaborations. Taken together, these findings further point to a democratizing influence of AlphaFold, allowing a broader set of institutions to participate more actively and more independently in high-impact protein research.

## Sample Construction

We examine the proportion of publications in leading journals for the top 500 universities over time. Publications are attributed to universities on the basis of the institutional affiliation of the authors. To identify top journals in protein research, we compile a list of 2,170 “biochemistry, genetics, and molecular biology” journals and 175 interdisciplinary journals publishing articles related to protein research. We rank the journals by their average SCImago Journal Rank (SJR) a journal reputation metric that accounts for citation counts of articles published in a given journal and the prestige of the journals where these citations appear. To account for annual fluctuations in SJR, we use the average SJR over the six-year period (2013–18) preceding AlphaFold’s release. There are 17 journals with an average SJR of 10 or above (see Table S2 for details), and we classify these journals as top journals.

We obtain data on all articles published in these 17 journals from OpenAlex (the successor to Microsoft Academic Graph). OpenAlex provides detailed article metadata including titles, abstracts, covered topics, citations, author affiliations, and the publishing journal. Each article is classified into one of three categories:

- *Structural biology*: if at least one author is a structural biologist or the paper covers any of 238 identified structural biology topics.
- *Non-structural protein*: if none of the authors is a structural biologist and the paper’s topics fall under one or more of 1,066 protein research topics outside structural biology.
- *Non-protein*: for all other micro-biology publications.

We rank universities based on their average total normalized citation scores (TNCS) between 2013 and 2018 in the category “biomedical and health sciences” (computed by Leiden University). The score for university *u* is defined as:

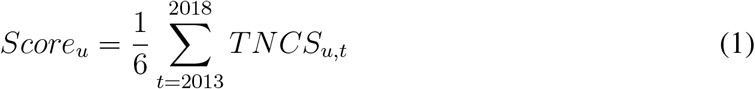

Where *TNCS*_*u,t*_ denotes the total normalized citation scores in year *t*. The top 12 universities are Harvard, Johns Hopkins, UCSF, Toronto, UPenn, Washington, UCL, Stanford, Duke, Michigan, UCLA and Oxford (Table S4).

Each author is assigned to a single university (highest ranked university for authors with multiple affiliations). The research output of each university *u* in year *t* is measured as the annual count of articles published in top journals where at least one author is affiliated with that university. We denote this output 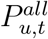 when we take into account all authors listed in a publication, and 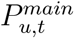 when we only account for the two main authors (listed first and last)^1^. The corresponding shares of top journal publications of university *u* in year *t*, which we refer to as “market share” is computed as follows:

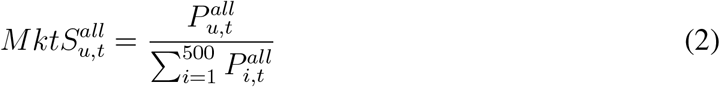

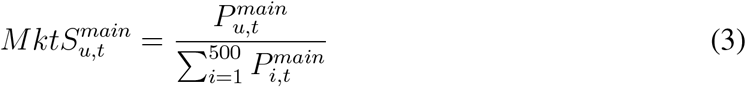

The cumulative density function (CDF) of 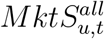 is quasi-monotone and concave (Figure 1). The top 10 universities generate 18% of all the top publications. The top 50 generate nearly half of all publications. The top 150 universities produce almost three times as many top publications as the next 350 universities (Figure 1(A)). Figure 1(B) shows that the distribution is similar if we only take into account the main authors (first and last authors) of each publication.

**Figure 1:**
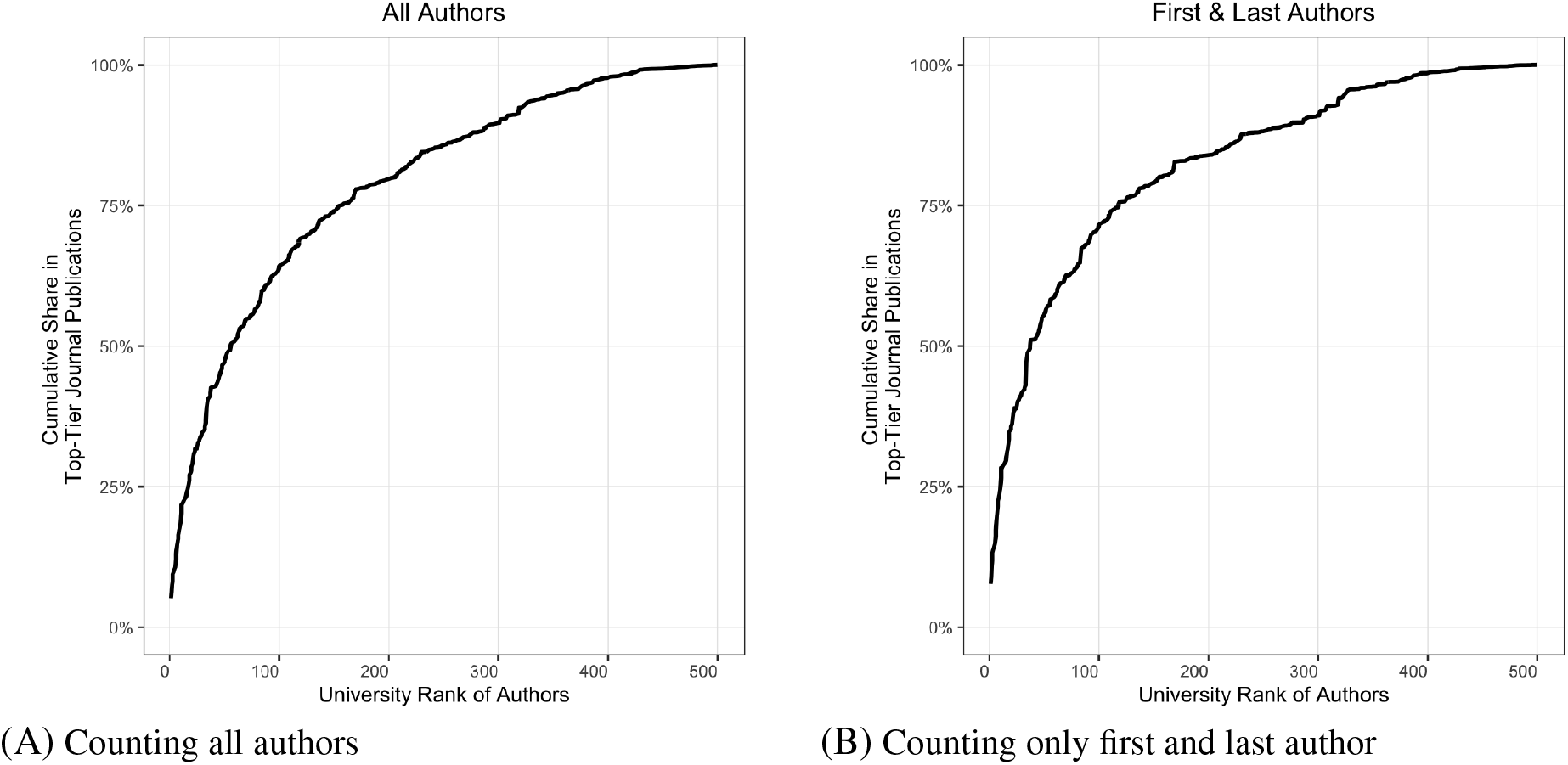
Market shares of top 500 universities before release of AlphaFold. University rankings are determined by the average *total normalized citation score* (TNCS) of each university in the “biomedical and health sciences” category between 2013 and 2018.

We rank universities by their market share of top journal publications from 2013 to 2018 and assign them to quartiles, each representing 25% of total output. The top quartile consists of 12 universities, followed by 36 in the second quartile, 88 in the third, and 364 in the bottom quartile. Geographically, top-quartile institutions are concentrated in the United States. The second quartile includes universities from the US and Europe, along with three in Australia. The third quartile is dominated by institutions in East Asia (notably China, Japan, and South Korea). The fourth quartile is more globally dispersed, with a concentration in Europe and East Asia and representation from South America, the Middle East, and Africa. See Figure S2 for a visual summary.

## Publication Rates around AlphaFold

### Changes in Aggregate Market Shares

We plot a time series of the aggregate market shares of each university quartile in Figure 2. In both panels 2(A) (structural biology publications) and 2(B) (non-structural biology, but within protein research), each quartile maintains a constant market share before the release of AlphaFold. The graphs also highlight three key milestones in the development of AlphaFold: (i) in 2018, AlphaFold 1 (AF1) wins the 13^*th*^ Critical Assessment of Protein Structure Prediction Competition (CASP); (ii) in 2020, AlphaFold 2 (AF2) wins CASP 14; and (iii) in 2021, millions of AlphaFold 2 predicted protein structures are made publicly available. Section S1.3 provides further details.

**Figure 2:**
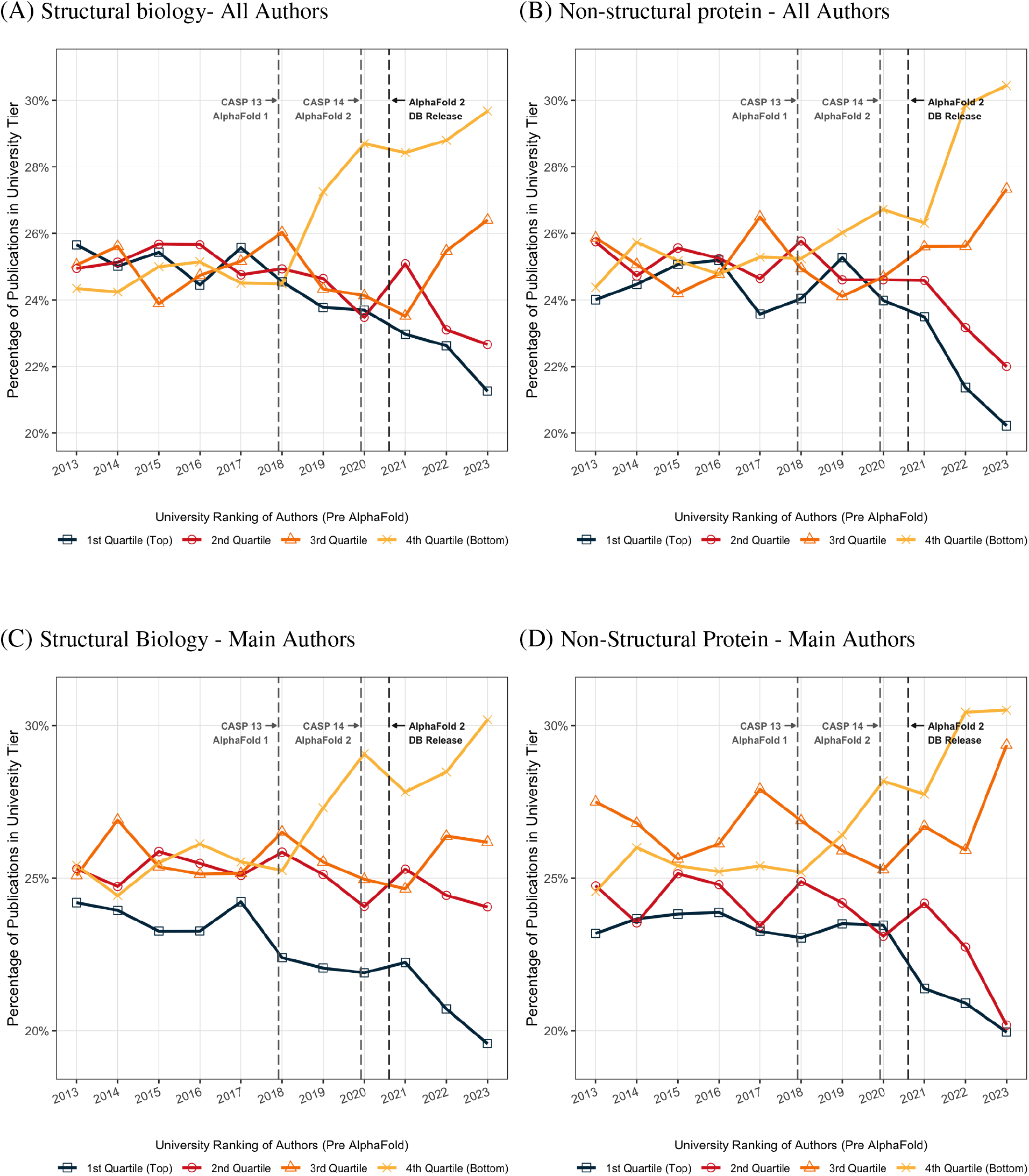
Protein publications in top journals by university quartiles. Figure illustrates the percentage of protein research publications in top journals (with an SJR ≥10 between 2013 and 2018, before the release of AlphaFold 1). The analysis is limited to authors affiliated with the top 500 universities, which are divided into four quartiles based on their share of publications in these top journals during the same period. Panel A and B show market share computed with all authors and panel C and D show market share computed with main authors only.

We observe a notable shift within structural biology publications after the release of AF1 (Figure 2(A)). Universities in the bottom quartile expanded their market share at the expense of those in the top two quartiles. In just two years, the market share of bottom-quartile universities rose from slightly above 24% to nearly 29%. This upward trend persisted with the release of AF2, and by 2023, the market share of the bottom quartile reached 30%, while the market share of the top two quartiles both fell to around 22%. A similar picture emerges when we compute market share only taking into account main -first and last-authors (Equation 3). Universities in the bottom quartile increased their share from 25% to over 30%. Meanwhile, the share of the top quartile dropped from 25% to 20% (Figures 2(C) and 2(D)).

The pattern observed in non-structural protein publications is also remarkable (panel 2(B)). No changes occur after AF1, but post AF2, there is a jump in top publications from institutions in the bottom quartile. The market share of universities in the bottom quartile increased from 26% to 30% in just two years. A similar, but less significant, increase is observed for universities in the third quartile. In contrast, institutions in the top two quartiles experienced a considerable reduction in market share, with the top quartile experiencing the largest decline.

We perform two sets of counterfactual analysis. We examine aggregate publication trends for i) publications in protein research journals with an average SJR between 1 and 10; ii) publications in our set of top-journals but excluding protein research. If the changes in the distribution of market shares observed in Figure 2 are driven by factors not related to AlphaFold, we would observe similar trends outside protein research (results shown in Figure 3) or outside the top journals (results shown in Figure S3). There is no pre-trend in either figure, and there is no effect post-AlphaFold either. The market shares remain the same in the period around AlphaFold’s release, showing that AlphaFold has only impacted protein research in the top journals specifically.

**Figure 3:**
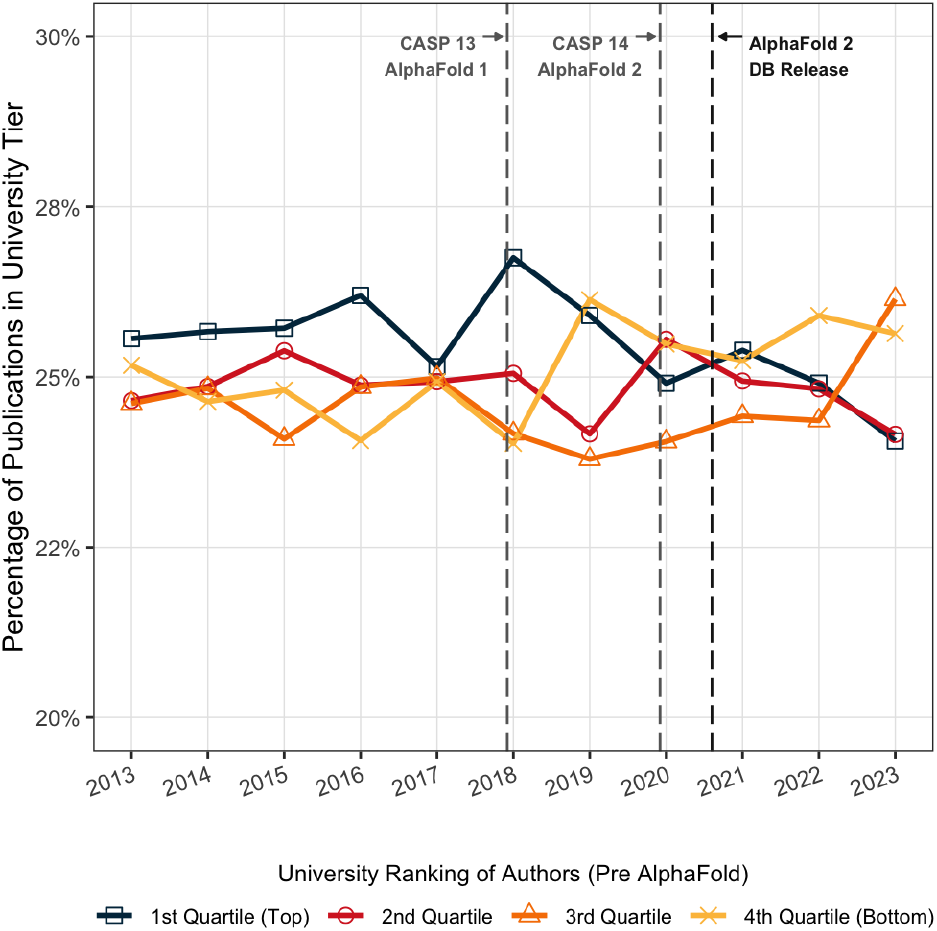
Publications in outside protein research in top journals by university quartiles. Figure illustrates the percentage of publications in microbiology excluding protein research in top journals (with an SJR ≥10 between 2013 and 2018, before the release of AlphaFold 1). The analysis is limited to authors affiliated with the top 500 universities, which are divided into four quartiles based on their share of publications in these top journals during the same period. University rankings are determined across all authors on a publication.

### Regression Analysis

To further test whether the changes in market shares observed in Figure 2 can be attributed to the introduction of AlphaFold, we employ a two-way fixed effects regression framework. Our analysis is conducted at the university level using a yearly panel of universities from 2013 to 2023. Each university is assigned to its pre-AlphaFold quartile as in Figure 2. This quartile-based approach allows us to estimate the average post-AlphaFold effect within each group, capturing shifts in publication dynamics while accounting for both time-invariant university characteristics and common yearly shocks. Specifically, for each university *u* in year *t*, we estimate the following equation:

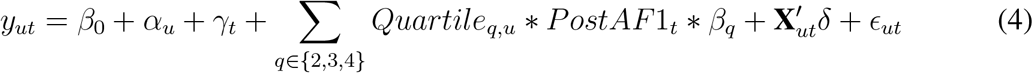

where *Quartile*_*q*_ is a binary variable equal to 1 if university *u* belongs to the quartile *u* (zero otherwise). *PostAF* 1 is a binary variable set to 1 for all years from 2019 onward, capturing the period after the introduction of AlphaFold. The vector **X**_*u,t*_ includes university-level control variables, with *δ* representing the corresponding coefficients. Fixed effects *α*_*u*_ control for time-invariant university characteristics, while *γ*_*t*_ capture common shocks affecting all universities in a given year.

We estimate the impact of AlphaFold on university research output using three variations of the outcome variable *y*_*ut*_. Each variation captures university publication activity in a different way. First, we define *y*_*ut*_ as the number of top-journal publications (in log) by all authors at university *u* in year *t*, to assess absolute changes in the number of publications. Second, we assess relative changes by defining *y*_*ut*_ as the university’s annual market share using equations 2 and 3, respectively.

We first estimate these models on the full sample of protein research publications. Panel 1(A) of Table 1 shows estimates for the three outcome variables. In each of these specifications, the interaction terms for the second, third, and fourth quartiles (relative to the top quartile) are positive and statistically significant at the 1% level. Ordinary least squares (OLS) estimates in model (1) indicate that the average fourth quartile university nearly doubled its number of yearly publications post–AlphaFold (increase by *e*^0.47^ *−* 1 = 0.60 from a 0.80 average). The effect is monotonic. The second and third quartiles exhibit the same upward trend, but with smaller magnitudes.

**Table 1:** Protein publications in top journals around AlphaFold. *PostAF1* is a binary variable set to one for papers published after the release of AlphaFold V1 in December 2018. The sample is limited to all protein research papers (both structural and non structural biology) published in top journals with an SJR score of 10 or higher and to authors affiliated with the top 500 universities prior to the release of AlphaFold 1. Universities are ranked based on their average TNCS in the period preceding the release of AlphaFold 1. The highest ranked universities placed in the top quartile (*University Tier = 1st Quartile*) are used as the reference group. Standard errors are clustered by university, and reported in parantheses. ***, **, and * denote significance at the 1%, 5%, and 10% level, respectively.

| Dependent Variables: | Log(1 + No. of Publications) | Market Share<br>(% Publications, All Authors) | Market Share<br>(% Publications, Main Authors) |
| --- | --- | --- | --- |
| Model: | (1) | (2) | (3) |
| University Tier = 2nd Quartile × PostAF1 | 0.19***<br>(0.02) | 0.06***<br>(0.00) | 0.10***<br>(0.00) |
| University Tier = 3rd Quartile × PostAF1 | 0.31***<br>(0.03) | 0.13**<br>(0.02) | 0.14**<br>(0.02) |
| University Tier = 4th Quartile × PostAF1 | 0.47***<br>(0.04) | 0.21***<br>(0.02) | 0.23***<br>(0.02) |
| Controls | ✓ | ✓ | ✓ |
| Year FE | ✓ | ✓ | ✓ |
| University FE | ✓ | ✓ | ✓ |
| Observations | 5,500 | 5,500 | 5,500 |
| Adjusted/Pseudo R<sup>2</sup> | 0.82 | 0.92 | 0.93 |

(B) Structural biology publications
| Dependent Variables: | Log(No. of Publications) | Market Share<br>(% Publications, All Authors) | Market Share<br>(% Publications, Main Authors) |
| --- | --- | --- | --- |
| Model: | (1) | (2) | (3) |
| University Tier = 2nd Quartile × PostAF1 | 0.23***<br>(0.02) | 0.13***<br>(0.00) | 0.15***<br>(0.00) |
| University Tier = 3rd Quartile × PostAF1 | 0.36***<br>(0.03) | 0.16***<br>(0.02) | 0.16***<br>(0.01) |
| University Tier = 4th Quartile × PostAF1 | 0.54***<br>(0.04) | 0.25***<br>(0.03) | 0.24***<br>(0.02) |
| Controls | ✓ | ✓ | ✓ |
| Year FE | ✓ | ✓ | ✓ |
| University FE | ✓ | ✓ | ✓ |
| Observations | 5,500 | 5,500 | 5,500 |
| Adjusted/Pseudo R<sup>2</sup> | 0.81 | 0.91 | 0.92 |

(C) Non-structural protein publications
| Dependent Variables: | Log(No. of Publications) | Market Share<br>(% Publications, All Authors) | Market Share<br>(% Publications, Main Authors) |
| --- | --- | --- | --- |
| Model: | (1) | (2) | (3) |
| University Tier = 2nd Quartile × PostAF1 | 0.28***<br>(0.03) | 0.09***<br>(0.01) | 0.12***<br>(0.01) |
| University Tier = 3rd Quartile × PostAF1 | 0.40***<br>(0.03) | 0.17***<br>(0.03) | 0.17***<br>(0.03) |
| University Tier = 4th Quartile × PostAF1 | 0.54***<br>(0.03) | 0.25***<br>(0.02) | 0.30***<br>(0.03) |
| Controls | ✓ | ✓ | ✓ |
| Year FE | ✓ | ✓ | ✓ |
| University FE | ✓ | ✓ | ✓ |
| Observations | 5,500 | 5,500 | 5,500 |
| Adjusted/Pseudo R<sup>2</sup> | 0.79 | 0.90 | 0.91 |

As university market shares are bounded between 0 and 1, models (2) and (3) use a generalized linear model with a logit link, which is essentially a logistic regression. This method transforms the probability of an event occurring into its log-odds ratio before relating it to a linear combination of predictor variables. Model (2) shows that following AlphaFold, fourth quartile universities show an average increase of 0.21 in the log-odds of the market share for top protein publications compared to top quartile peers. Similarly, second and third quartile universities experience an increase in log-odds, though with smaller magnitudes. Finally, estimates in model (3) confirm that our results hold even when market shares are based solely on the main authors.

We also conduct separate analyses for publications in structural and non-structural protein research (see Panels 1(B) and 1(C)). The estimated AlphaFold effect is equally strong across both domains, indicating that its impact reaches beyond structural biology alone. Note that the results reported here reflect average outcomes before and after AlphaFold’s release; annual trends are presented in Section S4.2.

### Changes in Research Practices

To explore the mechanisms underlying the observed shifts in market shares, we examine two key dimensions through which AlphaFold may have reshaped the protein research landscape and mitigated institutional disparities. First, we assess changes in research practices among university-affiliated authors using three metrics: *pivot behavior*, which captures the extent to which researchers shift their focus toward new or less familiar areas; *novelty*, which reflects the originality of a paper based on its engagement with rare or unconventional concepts; and *disruptiveness*, which indicates the degree to which a paper alters existing citation patterns by becoming a reference point independent of prior work.^2^ Second, we analyze shifts in collaboration patterns following the release of AlphaFold, focusing on co-authorship links between top- and lower-quartile universities to assess whether AlphaFold expanded access to resources and expertise traditionally concentrated in top-tier institutions.

Our analyses span both structural and non-structural protein publications in top journals across university quartiles. By examining changes in research orientation and collaborative networks, we assess two dimensions of scientific activity not captured by output-based metrics such as publication counts or market share. These dimensions are critical for understanding access to top-tier journals and may have played a role in the democratizing effects of AlphaFold documented above.

### Qualitative measures of research

Assessing the influence of AlphaFold on protein science requires looking beyond top journal publication counts alone. While such quantitative measures reflect changes in output, they do not fully capture potential shifts in the nature or impact of research. To complement our baseline analysis, we examine three qualitative metrics that reflect distinct dimensions of research activity at the author–paper level. These indicators offer insight into how AlphaFold may be reshaping scientific inquiry—enabling researchers to pursue new directions, introduce more original ideas, and contribute more meaningfully to ongoing debates in the field.^3^ Analyzing these metrics across university quartiles helps assess whether the rising presence of lower-quartile institutions in top journals is accompanied by changes in the character and potential impact of their research output.

The three metrics are defined as follows:

- **Pivot score**: This measure quantifies the extent to which a given paper by a researcher differs from their prior work based on the journals cited. It is calculated as 1 minus the cosine similarity between the distribution of journals referenced in the focal paper and those in the author’s previous publications over the past three years [22] (see detailed description of this measure in section S6). The pivot score ranges from 0 (no departure from prior words) to 1 (implying complete departure). A higher pivot score indicates a greater shift in research direction, which may be facilitated by AlphaFold’s provision of novel structural methodologies or predictions.
- **Novelty**: This metric evaluates originality in a paper by quantifying the rarity of concepts associated with its cited references, using OpenAlex topic classifications. As such, it follows an entity-based approach to estimating novelty [23–26], leveraging OpenAlex’s concept classification. For each reference, concepts with relevance scores above 0.5 are included, where the relevance score reflects centrality to the reference content. Novelty is then calculated as the average inverse of the works count for these concepts, with works count indicating prevalence in the broader literature (refer to Section S6 for a detailed description of this metric.). This approach denotes novelty because reliance on less common concepts signals that the paper builds on underrepresented ideas or unusual knowledge combinations, setting it apart from incremental advances in mainstream subject areas. Novelty matters in the context of AlphaFold because predicted 3D structures reduce the need for experiments requiring substantial resources, allowing researchers to study proteins that were previously inaccessible.
- **Disruptiveness**: This variable measures the degree to which a paper impacts subsequent research, based on citation patterns where future works cite the focal paper but not its predecessors [27, 28] (see Section S6 for a detailed description of this variable). A positive disruptiveness score indicates that the paper draws citations primarily to itself, implying it advances the field by reducing the prominence of earlier works. The provision of accurate protein structures by AlphaFold may facilitate this form of disruptive research, particularly for scientists at lower-tier institutions, thereby enhancing their influence in the field.

#### Changes in pivot behavior

Analysis of pivot scores reveals significant shifts in research direction across university quartiles following the introduction of AlphaFold. Prior to AlphaFold, researchers at top-quartile universities consistently exhibited higher pivot scores, indicating a greater propensity to make substantial shifts in their research direction. This pattern likely stems from their access to superior resources, funding, and collaborative networks, which facilitate exploration across diverse subfields of protein research. However, post-AlphaFold, this disparity narrows, with lower-quartile universities experiencing larger increases in pivot scores.

In structural biology, pivot scores increased across all university quartiles following AlphaFold, with larger gains observed in lower quartiles (Figure 4(A)). We quantify this effect using the regression specification in Equation 4, which estimates the differential impact of AlphaFold on pivot scores across quartiles while controlling for university fixed effects and time trends. Results from Model (1) in Table S8 show that, relative to the top quartile, bottom quartile universities experienced a 2.6% increase in pivot scores, third quartile universities a 1.7% increase, and second quartile universities a 1% increase, all highly statistically significant.

**Figure 4:**
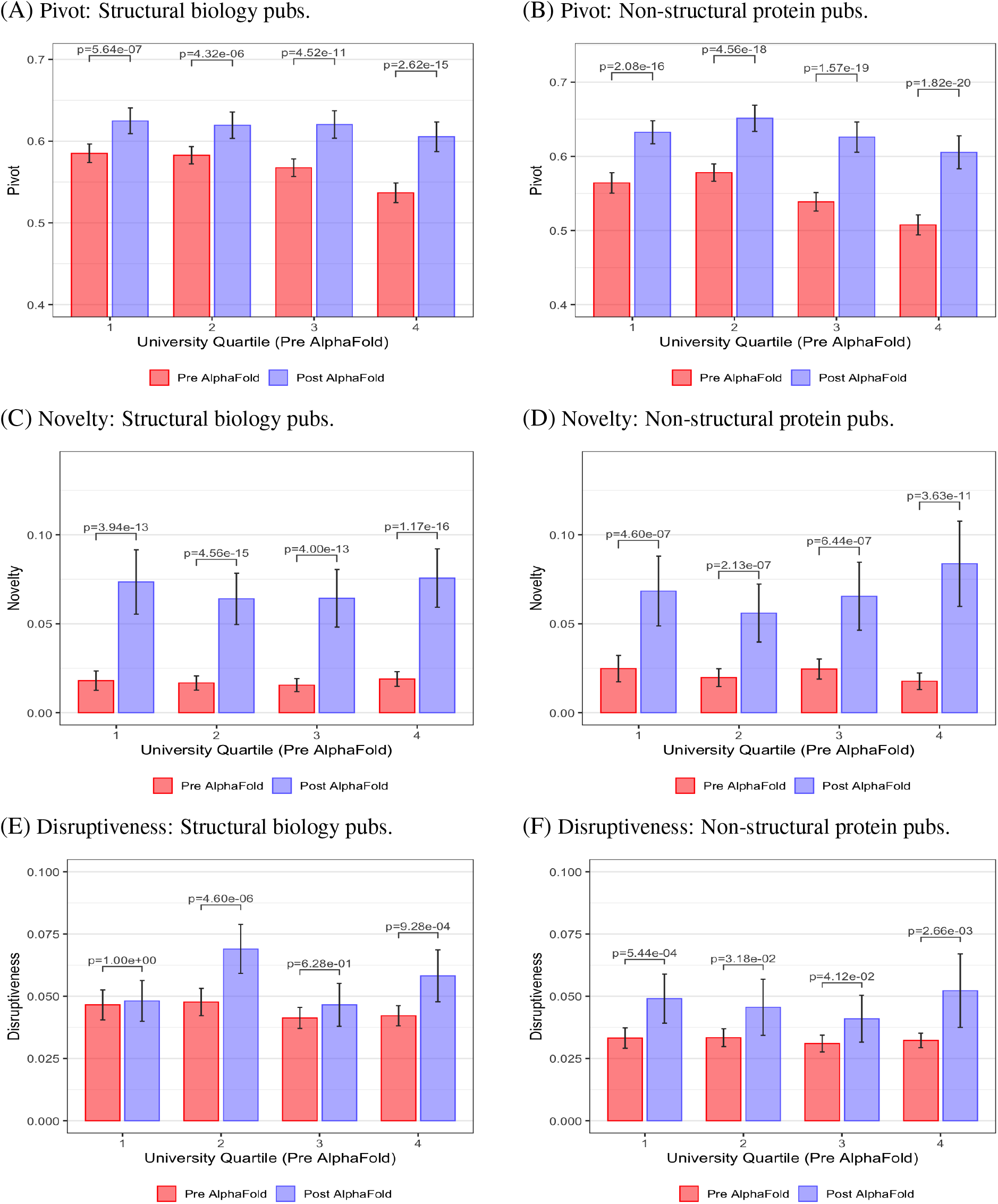
Research direction measures in top journal publications by university quartile. Figure shows pivot scores, novelty, and disruptiveness measures based on 3-year publication histories of authors in top journals (SJR ≥10). Panel (A) and (B) show pivot scores for structural biology and non-structural protein publications. Panel (C) and (D) show novelty measures. Panel (E) and (F) show disruptiveness measures. Universities are divided into four quartiles based on their aggregate share of publications in top journals during 2013-18.

We observe similar trends in non-structural protein research. Pivot scores increased across all university quartiles following AlphaFold, but with a more pronounced pattern favoring lower-tier institutions relative to those observed in structural biology (Figure 4(B)). We once again estimate this effect using Equation 4. Results from Model (4) in Table S8 show that, relative to the top quartile, second quartile universities experienced an increase of 3.4% in pivot scores and fourth quartile universities an increase of 3.3%, both highly statistically significant.

Taken together, these patterns suggest that AlphaFold, by offering a widely accessible tool for protein structure prediction, has helped lower barriers to entry in both structural and non-structural protein research. Researchers at lower-tier institutions appear better able to pursue new lines of inquiry and make more substantial shifts from their prior work (reflected in higher pivot scores) thereby narrowing some of the gap with top-quartile universities. These findings are consistent with prior work showing that technological shocks can broaden participation in science by reducing costs and enabling the exploration of ideas that are more novel or distant from researchers’ existing trajectories [29, 30].

#### Changes in novelty

Analysis of novelty scores reveals distinct patterns across university quartiles following the introduction of AlphaFold. These patterns likely stem from AlphaFold’s capacity to provide access to uncommon protein structures and predictions, allowing researchers to pursue hypotheses that were previously difficult to explore due to various constraints.

In structural biology, novelty scores increased across all university quartiles post-AlphaFold, with significant gains in lower quartiles (Figure 4(C)). Regression estimates using Equation 4 shown in Model (2) of Table S8 show that, relative to the top quartile, second quartile universities experienced a 2% increase, while third and fourth quartile universities both saw a 1.5% rise in novelty. These shifts underscore AlphaFold’s pivotal role in enabling researchers at resource-constrained institutions to address novel structural challenges, including the prediction of previously unresolved protein folds.

In the case of non-structural protein research, changes in novelty patterns are more pronounced, with novelty scores rising sharply for lower-quartile institutions (Figure 4(D)). The corresponding regression results presented in Model (5) of Table S8 indicate no significant effect for the second quartile but a substantial 1% (2.7%) increase for the third (fourth) quartile, relative to the top quartile. These results suggest that by leveraging AlphaFold-derived structural insights, researchers at lower-tier institutions are able to integrate accurate and new protein structures into non-structural domains such as drug discovery and functional genomics despite lacking experimental resources.^4^

#### Changes in disruptiveness

Disruptiveness, a citation-based metric, evaluates whether a paper consolidates (negative values) or disrupts (positive values) the existing citation network by attracting citations independently of its references. Our findings show that protein papers published in top journals post-AlphaFold have modestly redirected future studies away from prior literature. Due to citation lags, the full impact of AlphaFold on citation patterns may take years to emerge.

In structural biology, disruptiveness scores exhibit small but statistically significant increases across all university quartiles, with no clear trend favoring lower tiers (Figure 4(E)). Regression estimates from Model (3) in Table S8 indicate average annual increases of 1.3% for fourth-quartile universities, 1.2% for second-quartile, and 0.5% for third-quartile universities, relative to the top quartile. These modest gains suggest that AlphaFold supports incremental disruptions by providing accessible structural predictions that address challenges in traditional experimental methods, such as predicting complex protein folds previously difficult to resolve.

In non-structural protein research, disruptiveness effects are much weaker, with scores showing some overall increase (Figure 4(F)). Regression results in Model (6) of Table S8 reveal a small positive effect of 0.6% for fourth quartile universities and a small negative effect of -0.6% for third quartile universities, relative to the top quartile, both marginally significant at the 10% level. These subtle shifts indicate that AlphaFold’s structural insights have not yet significantly altered citation dynamics in non-structural protein domains. This may be due to the initial integration of structural predictions into studies aligning with existing research trajectories rather than immediately disrupting them. The limited impact could also reflect the time-lagged nature of citation-based metrics, where significant changes in research patterns often take years to fully emerge in the literature.

### Collaboration patterns

Collaboration represents another pathway through which AlphaFold, as a technological shock, may have reshaped the competitive dynamics of protein research publishing in top journals. Prior work shows that such shocks can influence collaboration patterns [29, 35, 36], which in turn have important implications for publication outcomes. High-impact research is increasingly produced through multi-university collaborations, which tend to involve institutions of comparable prestige [37–40]. Overcoming this tendency toward institutional homophily can be particularly beneficial for researchers at lower-ranked universities, who often see improved publication outcomes when collaborating with peers from top-tier institutions [29, 41–43]. Such collaborations offer access to advanced infrastructure, technical expertise, and greater visibility. Empirical evidence also indicates that the citation impact of multi-university teams tends to converge toward that of the more prestigious partner, effectively enhancing the scholarly reach of lower-tier institutions [40].

Accordingly, we examine whether AlphaFold increases the likelihood that researchers from lower-quartile institutions collaborate with peers from top-quartile universities—a dynamic that could disproportionately benefit the former and help account for their observed gains in top journal market share. Building on prior studies of team formation in science, we develop a model in which researchers form collaborations endogenously based on skill complementarities, subject to matching frictions [36, 44–47].

Our model posits that structural biologists are scarce and predominantly concentrated in top-quartile institutions. We conceptualize AlphaFold as a productivity shock for structural biologists, allowing them to participate in additional research projects and collaborations. Due to matching frictions, such as search costs that hinder the identification and evaluation of potential external partners, structural biologists tend to prioritize collaborations with colleagues within their own institution before seeking external partners. A key feature is that top institutions host a higher fraction of structural biologists relative to other protein researchers than in lower-quartile ones, as shown in our descriptive statistics (table S3). This disparity results in a smaller pool of potential internal collaborators per structural biologist. Consequently, top-quartile structural biologists deplete their internal collaboration opportunities more rapidly and are more likely to engage in external collaborations. These external partnerships disproportionately involve lower-quartile institutions, which lack sufficient internal structural biologists for collaboration. A formal exposition of this model is provided in Section S5.

To test this model, we estimate equation 4, using as the outcome variable the fraction of a university’s top journal publications in protein science that include at least one co-author from a top-quartile institution in a given year. In addition to the standard two-way interaction term *Quartile*_*q,u*_ *× PostAF* 1_*t*_, we include a triple interaction with the share of structural biologists in the university’s biology faculty for that year. This additional term is central to our empirical strategy, as the model predicts heterogeneous effects of AlphaFold on collaboration patterns depending on institutional structural biology capacity. Specifically, lower-quartile universities with few structural biologists are expected to show a larger decline in collaborations with top-quartile partners following the release of AlphaFold. The underlying mechanism is that AlphaFold may serve as a substitute for scarce structural biology expertise: researchers at resource-constrained institutions, who previously relied on top-tier collaborators to meet the technical demands of elite publishing, can now pursue more independent research, reducing the need for cross-tier collaborations even as top-quartile researchers become more productive.

Results are shown in Table S9, with the associated marginal effects in Figure 5. In the baseline specifications (models 1, 3, and 4), collaborations with top quartile authors decline across all lower quartiles following AlphaFold. The decline is steepest for fourth quartile universities, followed by third and second quartile institutions. This pattern reflects increased self reliance overall among researchers at lower tier universities, driven by AlphaFold provision of readily accessible protein structures.

**Figure 5:**
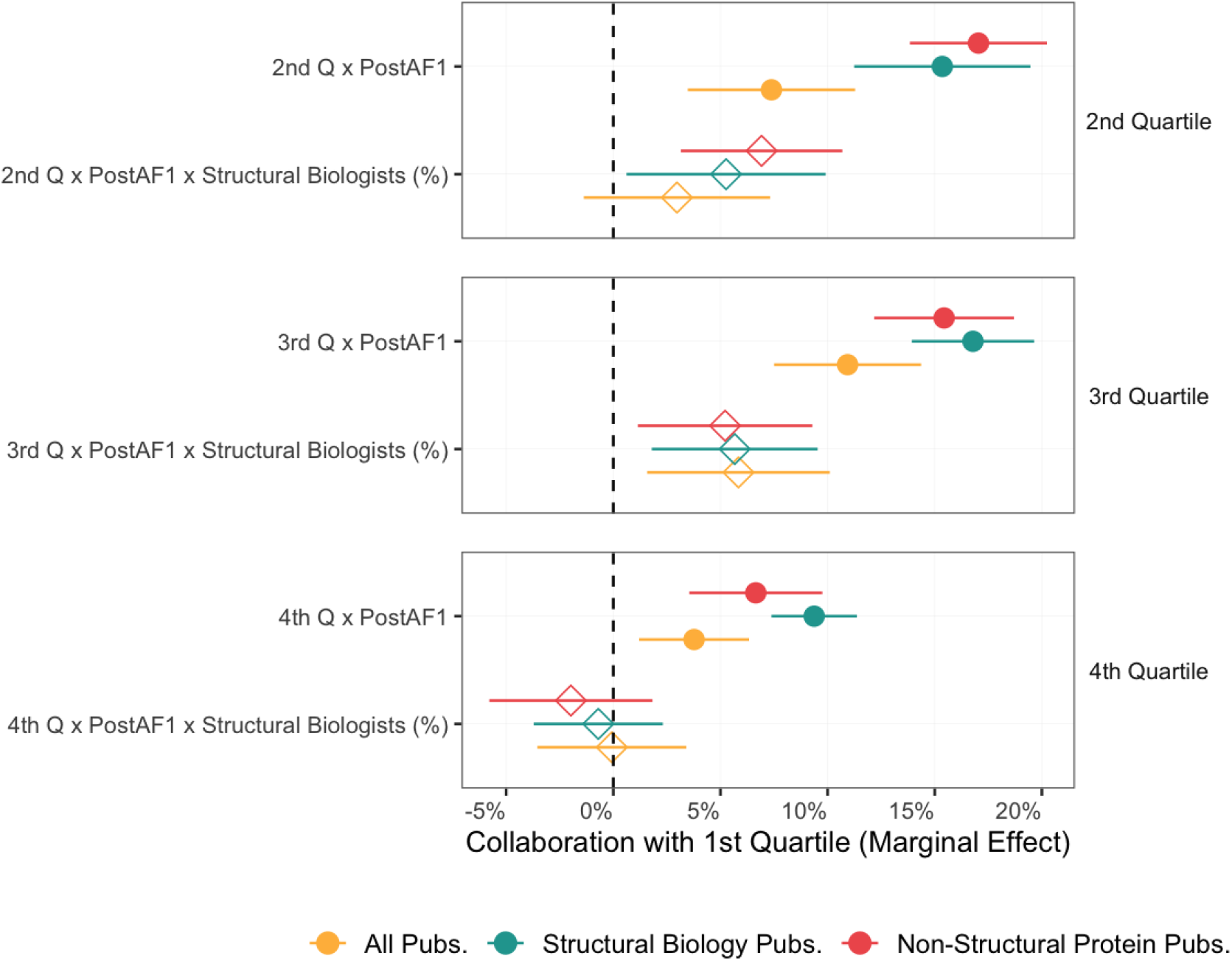
Collaborations with top-quartile universities in top journal publications around AlphaFold. This figure presents point estimates with 95% confidence intervals of the effect of AlphaFold on the frequency of collaboration between top-quartile authors and lower quartile authors for structural and non-structural protein publications. We specify a triple interaction structure to differentiate the effect of AlphaFold on top-quartile collaboration with universities rich in structural biologists, and those poor in structural biologists. The rest of the regression estimates can be found in Table S9. The coefficients displayed are a transformation of those shown in Table S9. For each triple interaction term (e.g., University Tier = 4th Quartile *×* PostAF1 × Structural Biologists (%)), the corresponding coefficients are multiplied by the average value of the corresponding variable and added to the double interaction (e.g., University Tier = 4th Quartile *×* PostAF1)

Models 2, 4, and 6 include the triple interaction with the share of structural biologists. In these specifications, the baseline post-AlphaFold effect — captured by the term *Quartile*_*q,u*_ *× PostAF* 1_*t*_ — is positive across lower-quartile universities, while the coefficients on the interaction term *Quartile*_*q,u*_ *× PostAF* 1_*t*_ *× Structural Biologists*(%) are negative and statistically significant. This pattern indicates heterogeneous effects: collaborations with top-quartile institutions increase post-AlphaFold primarily among lower-quartile universities with limited structural biology expertise, and this increase diminishes as the share of structural biologists rises. Although the per-institution collaboration gains are smaller among fourth-quartile universities with more structural capacity, the aggregate effect remains substantial given their large number. These findings are consistent with our model, which posits that the productivity boost from AlphaFold enables top-quartile structural biologists to take on more external collaborations — especially with institutions lacking in-house expertise — while institutions with stronger internal capabilities increasingly shift toward self-reliance in the post-AlphaFold period.

### Mediating effects of research quality and collaborations on publication outcomes

To understand the mechanisms underlying the observed increases in publication shares for lower-quartile universities following AlphaFold’s release, we next examine whether these gains are mediated by the changes in research quality and collaboration patterns documented in the preceding sections. Specifically, the improvements in pivot behavior, novelty, and disruptiveness, along with changes in the dependence on collaborations with top-quartile institutions, may explain how lower-tier protein researchers are achieving greater competitiveness in top journals. If these factors mediate AlphaFold’s impact, controlling for them should attenuate the estimated effects on publication outcomes. To test this hypothesis, we add controls for university-year average pivot scores, novelty, disruptiveness, and the share of collaborations with top-quartile authors to equation 4. The results appear in Tables S10 and S11 for structural biology and non-structural protein publications in top journals, respectively^5^.

Compared to the baseline effects documented in model 2 of Table 1(B), controlling for individual quality metrics in models 1–3 of Table S10 leaves the post-AlphaFold market share gains largely unchanged for the second and third quartiles in structural biology, with log-odds ratios shifting from 0.13 to 0.14 (a 1% increase in odds ratio) and from 0.16 to 0.15 (a 1% decrease), respectively.^6^ Strikingly, adding these controls substantially improves the estimated effect for fourth-quartile universities, with the log-odds ratio increasing from 0.25 to 0.37, corresponding to a 12.8% rise in the odds ratio. Non-structural protein research follows a similar pattern (models 1–3 of Table S11 relative to model 2 of Table 1(C)), with negligible shifts in log-odds for the second quartile (from 0.09 to 0.10, a 1.0% increase) and third quartile (from 0.17 to 0.16, a 1.0% decrease), but a pronounced rise for the fourth quartile, where the log-odds ratio increases from 0.24 to 0.34, raising the odds ratio by 10.5%. This increase in the estimated effect suggests that excluding quality metrics from the baseline model results in a downward bias. This bias arises from a tradeoff between publication quantity and quality, where increasing quality typically correlates with producing fewer publications (as researchers focus on higher-impact work). AlphaFold has improved both quantity and quality for lower-tier institutions, but the results suggest that the quality gains partially offset potential quantity increases due to this inherent tradeoff. In other words, if research quality had remained at pre-AlphaFold levels rather than improving, these institutions could still have improved their top publication volumes by reallocating effort from quality enhancements to producing more papers. The estimates remain stable even when all three quality metrics are included simultaneously in model 4, suggesting that they capture related aspects of the same underlying quality dimension.

Model 5 in both tables shows that controlling only for top-quartile collaborations leads to a modest increase in post-AlphaFold publication share gains across lower university quartiles. In structural biology, the log-odds ratios rise from 0.13 to 0.15 for the second quartile (a 2.0% increase in the odds ratio), 0.16 to 0.17 for the third quartile (a 1.0% increase), and 0.25 to 0.27 for the fourth quartile (a 2.0% increase). Non-structural protein research exhibits similar patterns, with log-odds increasing from 0.09 to 0.11 for the second quartile (a 2.0% increase), remaining stable at 0.17 for the third quartile, and rising from 0.24 to 0.25 for the fourth quartile (a 1.0% increase). These modest increases suggest that omitting collaboration with top-quartile authors from the baseline model introduces a small downward bias. This bias arises because such collaborations are positively associated with top publication outcomes. However, lower-quartile researchers, particularly those at universities with more structural biology expertise, engage in fewer such collaborations post-AlphaFold, as evidenced in Table S9. By controlling for collaboration, we mitigate this bias, uncovering a slightly larger underlying effect on market shares.

Model 6 adds controls for both research quality and collaborations with top-quartile authors. Relative to the preceding models, including these controls does not change the estimated post-AlphaFold publication share gains by much, though they remain higher than in the baseline specification.

Overall, the estimated impact of AlphaFold on top protein publications becomes more pronounced after accounting for concurrent changes in research quality and collaboration patterns. However, these results warrant cautious interpretation, as metrics such as pivot behavior, novelty, and disruptiveness are likely endogenous to collaboration dynamics. Shifts in co-authorship can expose researchers to new ideas and domains, potentially influencing both the direction of their work and the structure of citation networks [45, 48].

## Discussion

The persistent concentration of scientific output within elite institutions has long been attributed to unequal access to resources, infrastructure, and reputational capital rather than differences in individual researcher ability. Our findings provide empirical support for the idea that AI can disrupt these structural advantages. The introduction of AlphaFold represents an exogenous technological shock that reduced the skill and resource intensity historically required for cutting-edge protein research, particularly in structural biology. This shock has enabled researchers at less prestigious institutions to increase their presence in top-tier journals, suggesting that AI can function as a scientific equalizer. Notably, the effects are both quantitative and qualitative: lower-quartile universities not only increased their publication volumes and market shares in top-tier journals but also exhibited significant improvements in research quality relative to elite institutions, thereby narrowing preexisting institutional gaps in innovation and impact within protein science.

These results align with prior work on the redistributive effects of technological shocks in science. Prior studies have shown that when experimental costs fall or computational tools become more accessible, the institutional distribution of research can shift [30, 35, 49]. Yet, whereas such shocks manifest gradually and are limited in scope, AlphaFold’s effects have been rapid and large, possibly pointing to a different mechanism: one linked to the general purpose and open-access nature of modern AI tools. Intriguingly, our robustness checks reveal that controlling for concurrent improvements in research quality and shifts in collaboration patterns actually strengthens the estimated publication gains for lower-quartile universities. This indicates that improvements in research quality and shifts in cross-university collaborations only partially account for the observed changes in top journal market shares. Additional factors, such as AlphaFold’s capacity to facilitate independent, high-impact research by bypassing the need for proprietary infrastructure, are likely key drivers of the enhanced competitiveness of resource-constrained institutions.

Our findings also contribute to long-standing debates about the Matthew effect in science. Elite universities have historically leveraged cumulative advantages in funding, editorial control, and collaboration networks to reinforce their dominance [8, 9]. The changes we observe after AlphaFold suggest that such feedback loops can be interrupted, particularly when new tools alter the balance of what it takes to produce top quality research. For instance, lower-quartile researchers became less reliant on collaborations with top-quartile institutions to access top journals, as evidenced by an overall decrease in their propensity for such partnerships. This reduced dependence highlights how AlphaFold democratized access to critical structural insights, allowing lower-tier institutions to pursue ambitious projects autonomously. Importantly, the accessibility of AlphaFold played a central role: unlike many scientific breakthroughs constrained by proprietary datasets or infrastructure, AlphaFold provided a freely available computational model and a comprehensive, open-access protein structure database, effectively diluting traditional barriers to entry in structural biology research.

Whether future AI innovations will follow AlphaFold’s trajectory remains to be seen. As AI tools become more complex and expensive to train, the risk increases that control over their development and deployment will centralize within a small number of private firms. AlphaFold itself was developed by Google DeepMind and made freely available, but future iterations of protein prediction models may be gated by proprietary data, computational scale, or commercial licensing. This raises concerns that the current moment of democratization could be temporary, replaced by new hierarchies tied to access to AI infrastructure.

Furthermore, while AlphaFold reduces the cost of structure prediction, many downstream research activities such as experimental validation and drug discovery remain capital intensive and institutionally skewed, often requiring access to advanced facilities, proprietary datasets, or interdisciplinary expertise disproportionately available at elite institutions. As such, the increased publication volume and improved research quality we document for lower-quartile universities may not immediately lead to comparable advances in practical applications, such as faster development of new therapies or more equitable access to research funding and infrastructure.

Future research should examine whether the rebalancing of top publication shares observed post-AlphaFold represents a lasting shift, translating into more durable changes in scientific influence, collaboration networks, and funding allocation. It will also be important to assess whether these democratizing effects extend beyond protein science to other domains such as genomics, materials science, and climate modeling where AI tools may similarly empower underrepresented institutions or, conversely, introduce new forms of exclusion.

Recent developments surrounding AlphaFold 3 highlight the fragility of such gains. It was initially released with restricted access in May 2024, which prompted widespread calls from the scientific community for greater access. It was ultimately made open access in November 2024. This episode underscores the importance of longitudinal studies on the evolving role of AI in research ecosystems, particularly regarding how access and governance decisions shape equitable innovation.

As AI continues to transform the landscape of scientific discovery, global policy responses are accelerating. The year 2025 has already seen major initiatives, including the U.S. AI Action Plan and international efforts emphasizing equitable access to AI technologies. In this context, a deeper understanding of AI’s potential to either reinforce or reduce institutional hierarchies will be essential for designing inclusive science and innovation policies that broaden participation and enhance societal benefit.

## Acknowledgments

We are grateful to Pierre Azoulay, Kyle Myers, Jacqueline Lane, Fabrizio Dell’Acqua, Hans Hvide, Savvas Savvides, Rob van der Kant, Emiel Michiels, and Lucrezia Vittoria Viti for their invaluable feedback and support. We also thank seminar participants at Harvard University, University of Bergen, and University of Illinois for useful comments.

## Supplementary Information

### S1 Literature & Background

#### S1.1 Market Concentration in Science

Scientific research is highly concentrated, with the top 1% of scientists obtaining 20% of the citations [1] and 50% of funding [2, 3]. The pros and cons of this concentration are actively debated [4]. Advocates for a more even distribution argue that concentrating funds stifles scientists’ creativity.For example, superstar scientists may act as gatekeepers and hinder the emergence of new paradigms [5]. Similarly, in grant reviews, it has been found that homophily among senior scientists can penalize the more novel research proposals [6–8]. In contrast, critics point to economies of scale and efficient resource use by more productive scientists.

Research in this area has focused on explaining this concentration. A prevailing explanation is that scientific concentration arises partly because of the winner take all reward system in science: renowned scientists receive a disproportionate share of funding and credit for their research. This boosts their productivity and fame, thus further entrenching their position at the top. A phenomenon known as the “Matthew effect” [9]. This theory has been empirically tested in various contexts [3, 10, 11].

Related to our work are papers examining the effect of external shocks on competition in science. The hiring of stars in lower-ranked institutions boosts the productivity of future recruits without affecting existing researchers [12]. The deaths of stars’ scientists negatively impact their collaborators but benefit others in the same field [5, 13]. The influx of Soviet mathematicians decreased career opportunities for prominent American mathematicians post-USSR [14]. Bitnet improved collaboration and publication results for mid-tier researchers located near top scientists [15]. The introduction of CRISPR, a novel gene editing tool, first improved the prospect of institutions that were already at the cutting edge of genetic research [16]. The free availability of Microsoft’s Kinect democratized motion sensing technology, increasing the competitiveness of outsider researchers and the diversity of ideas [17]. The reduced costs for accessing genetically modified mice led to new researchers entering the field and to more diverse research being produced [18].

#### S1.2 Structural Biology

Structural biology is a branch of protein research, alongside genomics, drug discovery etc. Its primary objective is to identify the three-dimensional shapes of proteins. Knowing the structure of proteins is key to understand how they interact with various molecules [19] and is fundamental for drug development [20]. Before 2018, most protein structures were determined by X-ray crystallography [21]. The cost of this technique varies between $250,000 for soluble human proteins (25% success rate) and $1,000,000 for membrane proteins (10% success rate) [22].

Proteins are crystallized by preparing a concentrated protein solution to induce crystal formation. This crystallization process is complex and highly variable, often involving extensive trial and error with conditions such as temperature, pH, and specific chemical additives [23]. Once suitable crystals are obtained, they are exposed to high-intensity X-ray beams at a synchrotron, which produces a diffraction pattern that reveals the atomic arrangement. The diffraction data are transformed into an electron density map that shows the atomic positions within the protein. The scientists then test and refine their model to align with the data. This iterative process can take months, requiring multiple adjustments to accurately capture structural details [24].

An alternative to X-ray crystallography is cryo-electron microscopy (cryo-EM). Cryo-EM freezes proteins at cryogenic temperatures, maintaining their natural structures. An electron beam captures 2D images that are computationally assembled into a 3D model. Cryo-EM is popular for resolving large, complex, and membrane protein structures [25, 26].

#### S1.3 The Advent of AlphaFold

AlphaFold (AF) was developed by London-based DeepMind AI lab (part of Google). Their first model used a neural network trained on experimentally determined 3D structures of proteins from the Protein Data Bank. The DeepMind team was not the first to attempt protein structure prediction using deep learning but was the first to succeed. They used a much larger, and more computationally efficient neural network than previously [27]. AF processed amino acid relationships and inferred structural data faster and more accurately than traditional methods [28], and won the 13^*th*^ Critical Assessment of Structure Prediction (CASP) competition.

The results of CASP13 validated the deep learning approach after years of limited success. It sparked the development of various deep learning models to predict structures. The associated research paper was subsequently published in the journal Nature in January 2020.

AF precision was still insufficient for reliable predictions of complex proteins, and in 2020, AlphaFold 2 (AF2) was introduced. It combined evolutionary data, structural templates, and a revised deep learning infrastructure that featured an attention mechanism. Compared to AF1, AF2 better understood the spatial dependencies between amino acids and achieved precision comparable to experimental methods [29]. On 30 November 2020, it won the CASP 14 competition by a record margin.^7^

On 15 July 2021, the paper introducing AlphaFold 2 was published in Nature as an advance access article [30]. The publication came with an open source software and a searchable database with predicted structures for more than 214 million proteins, covering most known natural sequences, including the entire human proteome. In contrast, pre-AF, only 17% of the human protein structures were known [31]. The paper received more than 20,000 citations in two years, and its main authors won the 2024 Nobel Prize in Chemistry.

Since the release of AF2, several deep learning tools emerged to address structural protein challenges (Table S1 provides examples), and on 8 May 2024, DeepMind announced the release of an enhanced version (AlphaFold 3). AF3 is capable of predicting highly complex protein structures and protein-protein interactions. The development of AlphaFold and other models is summarized in figure S1.

### S1.4 Role of AlphaFold in Structural Biology

AlphaFold can serve both as a complement and a substitute to experimental methods in structural biology. AlphaFold predictions can be used either to bypass experimental work entirely or to automate some aspect of the experimental process.

As a substitute, AlphaFold enables researchers to obtain predicted structures without costly experiments [31].The availability of predicted structures surged with the AF2 database, allowing easy access to structures without needing to run a model like AlphaFold. However, exclusive reliance on AlphaFold without experimental validation may jeopardize accuracy. AlphaFold 2 can encounter difficulties with novel or large protein complexes in particular, leading to inaccurate predictions even when prediction confidence intervals are low [32].

As a complement, AlphaFold makes experimental structure discovery faster and cheaper. AlphaFold predictions provide more complete starting hypotheses to build models that better fit the experimental data. AlphaFold predictions also enable faster and cheaper trial and error over large sets of potential structures. More iterations lead to finer interpretations of experimental data, thus improving structure determination [33].

AlphaFold does not only improve productivity for existing tasks, it also enables the study of larger protein systems. For instance, the nuclear pore complex, crucial for nuclear transport, presented considerable modeling challenges. A detailed structure was constructed using various advanced experimental techniques to integrate individual segments into a unified model [34]. Subsequently, this model was refined using predictions derived from AlphaFold[35]. Such advanced models would be unattainable without AlphaFold.

### S2 Data and Sample Construction

Our main data source is OpenAlex. OpenAlex was developed by OurResearch, a nonprofit organization, to facilitate access to the Microsoft Academic Graph after Microsoft discontinued the project in 2021.^8^ Each publication is assigned a unique identifier, and its metadata include the title of the article, the abstract, citation counts, the list of authors with their institutional affiliations and departmental affiliation, and the range of topics covered.

#### S2.1 Journal Selection

We select journals in ‘biochemistry, genetics, molecular biology’ (2,170 journals) and ‘Multidisciplinary’ (175 journals), then label those with an average SJR greater than 10 as ‘top journals’ (17 in total, listed in Table S2). We split the articles published in these journals into three groups labeled as follows: i) ‘Structural biology’, ii) ‘Non-structural protein’, and iii) ‘Non-protein.’ A paper is classified as structural biology if it has an author who is a structural biologist or if the paper list of topics includes one of the 238 topics we identify as structural biology. If a paper does not satisfy this condition but has one of the 1,066 topics of protein research, then it falls in the second group. All other papers are assigned to the third group. Figure S4 shows category trends over time. In 2013, “non-protein” research accounted for 50% of publications, with protein-related studies split equally (25% each). By 2024, protein-related research dropped to 20% (10% each for structural and non-structural), while non-protein research grew. Overall publications declined from 4,000 in 2013 to 2,800 in 2024, independent of COVID-19 research.

We measure the quality of the journal in which the research is published with the SCImago Journal Rank (*SJR*). SCImago is an online platform that provides a comprehensive list of academic journals in various disciplines, together with statistics such as citation counts and the proportion of female authors. To generate its SJR, SCImago employs an algorithm that uses both the number of citations of articles published in a given journal and the prestige of the journals where the citations appear. As the SJR of each journal fluctuates from year to year, we rank the journals using the average SJR during the six-year period preceding the release of AlphaFold (i.e., 2013–2018).

#### S2.2 University Ranking

The Centre for Science and Technology Studies (CWTS) at Leiden University produces the *Total Normalized Citation Score* (TNCS) of about one thousand universities around the world. We compute the average TNCS score of each university between 2013 and 2018, in the category “biomedical and health sciences”. TNCS is computed as follows. First, each publication is assigned to a biomedical subfield. Then, the citation count for each publication is compared to the average in the subfield in that year, resulting in a normalized citation score, where 1 represents the overall average. The Mean Normalized Citation Score (MNCS) is then calculated by averaging these normalized scores across all the publications in the subfield. Finally, the TNCS is obtained by multiplying the MNCS by the total number of publications, providing a size-dependent indicator of the university’s overall citations. This method thus accounts for differences in citation practices between subfields and across years.We match authors to universities based on their affiliation. In our data some authors are not affiliated with universities, but instead with research centers, private companies or hospitals. These researchers are dropped from the data.

We compute the yearly publication count of university *u* as follows:

Let

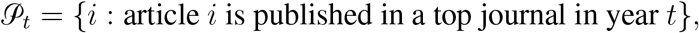

and for each article *i*, let

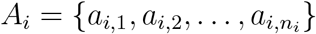

denote its ordered set of authors. Denote by *u*(*a*) the university assigned to author *a* (i.e., the highest ranked among any multiple affiliations).

Then, the publication counts are defined as follows:

1. All authors count:

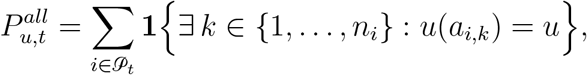
2. Main authors count:

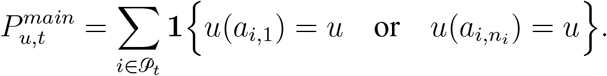

These definitions ensure that each article contributes at most one count per university, regardless of the number of affiliated authors.

For robustness, we also construct a measure of market share that only takes into account the two main contributors on a publication (first and last author in the author list).

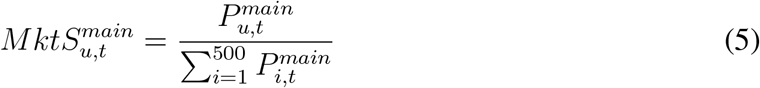

Market concentration is slightly stronger when we only account for the main authors. The top 50 universities then generate more than half of all top publications.

#### S2.3 Additional Data

High-performance computing (HPC) plays a pivotal role in modern protein research, enabling complex simulations, large-scale data analyses, and computationally intensive methods such as those employed by AlphaFold. Access to substantial HPC resources is crucial for enhancing researchers’ productivity and enabling the exploration of a broader range of research topics[37]. To empirically assess the influence of HPC on protein research outputs, we use data from Top500.org as a proxy for the computational power available at academic institutions. Since 1993, Top500.org has systematically tracked HPC facilities at universities worldwide on a biannual basis, providing detailed metrics on total processing power (quantified by the number of processor cores) and computational speed (measured in petaflops per second). This dataset allows us to assess the link between a university’s HPC capabilities and its structural biology output. We compare research volume and quality against available computational power over time. This analysis reveals whether advanced HPC drives high-quality protein research, especially in studies requiring structural biology expertise.

#### S2.4 Descriptive statistics

A range of descriptive statistics is presented in table S3. They highlight significant disparities in research production across university quartiles prior to AlphaFold. Universities in the first quartile produce more in both structural and nonstructural protein research. They average 8.58 publications on structural biology (per university annually), compared to 2.39, 1.75, and 0.8 for the second, third, and fourth quartiles, respectively. The numbers are similar in non-structural protein research. It follows that first-quartile universities also hold a larger market share, both in terms of the share of publication and the share of authorship. The average first quartile university produces 0.37% of all the top structural biology publications, in contrast to 0.1%, % and 0.03% for the second, third and fourth quartile, respectively. The numbers are similar for non-structural protein publication and authorship share.

Table S3 indicates that top-quartile universities have more resources at their disposal. The average faculty size is nearly double that of the second quartile and more than triple that of the fourth quartile. Access to HPC resources varies greatly. The first quartile universities have a higher rate of HPC access (50.9% versus 9% of fourth quartile universities), and when they do, have more computing power (351.11 petaFlops versus 54 petaFlops).

In terms of geographical distribution, American universities are the most prevalent in quartile 1, European universities peak in quartile 2, and Asian universities in quartile 3. Additionally, the top-quartile universities have more researchers affiliated with industry but fewer with governmental labs. There are few differences between quartiles in other dimensions. For example, the racial and gender distribution of the average university is similar across quartiles: around one third female, half Asian, one quarter white.

### S3 Empirical Design

We split the publication distribution into four quartiles, each representing approximately 25% of all top journal publications and categorize universities accordingly. The first quartile includes the top 12 universities. The second quartile includes the next 36 universities, the third quartile the next 88, and the fourth quartile the next 364.

OpenAlex does not provide demographic indications for authors. We determine author ethnicity using the rethnicity package in R. This package predicts ethnicity from first names using probabilistic models trained on demographic data. Gender is identified by the genderizeR package in R, using first names and a large online database. Researchers in departments specializing in structural or computational biology or similar fields are classified as structural biologists.

In the main regression specifications, we include as control all of the variables mentioned in Table S3. to control for changes in faculty composition. The dependent variable *y*_*ut*_ is one of three variables:

1. The log number of publications in top journals from university *u* at time *t*: 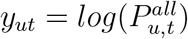
2. The market share of university *u* at time *t*: 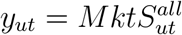
3. The market share of university *u* at time *t* taking into account only main authors is: 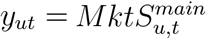

### S4 Additional Results

#### S4.1 AlphaFold Adoption

We approximate the use of AlphaFold or other equivalent AI systems at the publication level. We classify a publication as using AlphaFold if i) it explicitly mentions AlphaFold or an analogous AI tool in their abstract, or ii) it cites the original AlphaFold paper [38], the main paper that introduced AlphaFold. Note that this method for identifying AlphaFold usage is neither a lower-bound nor an upper bound of real adoption: one paper might be mentioning or citing AlphaFold without using it, while another paper might be using AlphaFold without mentioning it or citing it. With that being said, we do expect our measure to underestimate the use of AI more than it overestimates it: We can only search for AlphaFold mentions in the abstract, and we expect that many publications mention AlphaFold only in the body of the text.

Figure S5 shows the proportion of publications using AlphaFold in both structural and non-structural protein publications, as well as in different subfields of structural biology. We observe a relatively modest adoption in 2018, which remains stable for three years, before a large increase follows the release of the AF2 database. Subsequently, we see a rapid adoption in structural biology, reaching 30% in 2023, as shown in figure S5(A). The results are similar for nonstructural protein research, but the magnitudes are smaller. If we restrict the sample to papers on protein structure prediction and macromolecular crystallography, the adoption rate reaches 60% (Figure S5(B)).

Adoption rates also vary between university quartiles. The panel S7(A) shows the AlphaFold share of publications for each university quartile. In structural biology, the two lower quartiles produce the majority of AlphaFold articles (36% and 31%, respectively). The picture is more nuanced in nonstructural protein research. Panel S7(B) indicates that all quartiles had a similar AlphaFold paper output between 2020 and 2022, with a stronger adoption by the bottom quartile pre-2020. In the last year of our sample we see a shift where bottom quartile shares reach 30% (quartile three) and 34% (bottom quartile), while the shares of quartile one and quartile fall to 16% and 20% respectively.

#### S4.2 Dynamic estimates

We complement the fixed effects regressions from equation 4 with yearly cross-sectional estimates using identical controls. Figures S8 and S9 reveal distinct temporal patterns. Post-AlphaFold, publication volumes in the bottom three quartiles increase relative to the top quartile, with a pronounced 2020 spike followed by a 2021 decline and subsequent annual growth. Effects are stronger and more precise in structural biology than broader protein research.

Market share estimates exhibit similar dynamics with greater noise and wider confidence intervals. The 2020 spike coincides with the COVID-19 pandemic, complicating interpretation. However, estimates from 2021 onward align with the gradual AlphaFold adoption documented in section S4.1.

#### S5 Collaboration dynamics

We formalize how an increase in project capacity shifts collaborations of top-tier 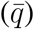 structural biologists toward lower-quartile partners. We distinguish between partners from sbio-poor schools (*q*_*p*_) and sbio-rich schools (*q*_*r*_), and introduce a friction that makes the latter less available for collaboration.

Let:

- 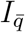be the set of top-quartile structural biologists, 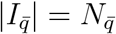.
- 𝒥 be the universe of potential collaboration slots. Each slot *j* is associated with a partner type 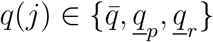.
- *X*_*j*_ be the random payoff of slot *j* ∈ 𝒥, drawn from a distribution *F*_*q*(*j*)_ with mean *µ*_*q*(*j*)_.

We assume payoffs are ordered by partner tier: 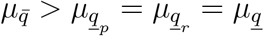.

- *B* be the number of projects a top-quartile biologist can engage in.
- *M* be the number of potential slots a 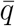 researcher can sample from each partner type.

We assume researchers at sbio-rich schools (*q*_*r*_) face internal collaboration demands. A fraction *γ* ∈ (0, 1) of their collaboration capacity is reserved for local projects, making them unavailable for external collaboration with 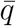 researchers. This reduces the effective pool of available partners from sbio-rich schools.

For each biologist *i*, the set of available collaboration slots is:

- 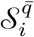: A random sample of *M* slots from the 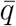 pool.
- 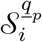: A random sample of *M* slots from the *<u>q</u>*_*p*_ pool.
- 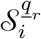 : A random sample of (1 *− γ*)*M* slots from the *<u>q</u>*_*r*_ pool.

Let 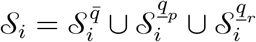 be the total set of projects available to researcher *i*.

Each structural biologist *i* observes realized payoffs from their available slots and selects the top *B* projects:

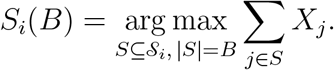

Define the share of collaborations with partner type *q*:

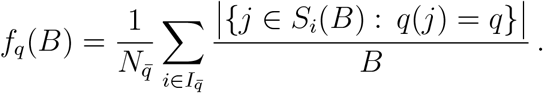

Let 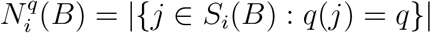 be the number of collaborations of type *q* for researcher *i*. The change in the share *f*_*q*_(*B*) as *B* increases to *B* + 1 depends on the type of the newly added project, *j*^*∗*^.

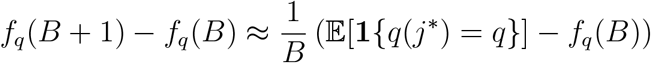

The term is positive if the probability of the marginal project being of type *q* is greater than the current share of type-*q* projects.

As *B* increases, the highest-payoff projects (mostly with 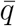 partners) are selected first. The marginal project *j*^*∗*^ is therefore increasingly likely to be a lower-quartile collaboration. Since the pool of available *q*_*p*_ partners is larger than the pool of available *q*_*r*_ partners (*M >* (1*−γ*)*M*), the probability that the marginal project is with a *q* _*p*_ partner is greater. Thus we have:

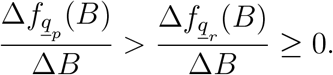

### S6 Qualitative changes

We employ three metrics to quantify changes in research direction: pivot score, novelty, and disruptiveness. These measures capture different dimensions of scientific research orientation and allow us to assess how AlphaFold affected research directions across university quartiles.

#### S6.1 Pivot Score

The pivot score measures the extent to which a researcher’s new work diverges from their previous research portfolio. We compute this score using cosine similarity between the distribution of journals cited in a new paper and those cited across the researcher’s prior work.

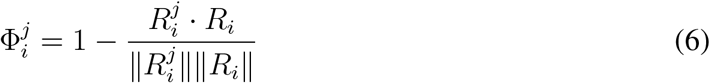

where 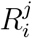 is a vector representing citation frequencies to journals in focal paper *j* by researcher *i*, and *R*_*i*_ aggregates citation frequencies across all prior papers by researcher *i*. The score ranges from 0 (no departure from prior work) to 1 (complete shift in research direction).

#### S6.2 Novelty

Novelty quantifies a paper’s originality by assessing the rarity of concepts in its references, using OpenAlex concept classifications. Each concept has a relevance score (0-1) indicating its centrality to the paper and a works count reflecting its prevalence across the literature.^9^

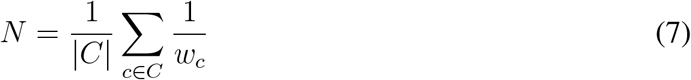

where *C* is the set of unique concept IDs from the paper’s references with relevance score *>* 0.5, and *w*_*c*_ is the works count of concept *c*. The 0.5 threshold ensures only concepts meaningfully connected to the references contribute to the score. The inverse weighting 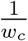 assigns greater importance to rarer concepts, so papers referencing niche topics with low works counts (e.g., 50 papers) receive higher novelty scores than those relying on widely studied topics (e.g., 10,000 papers). For example, a paper on neural networks might have high relevance for “machine learning” (relevance = 0.9) but low relevance for “agriculture” (relevance = 0.2), with only the former contributing to the novelty calculation.

#### S6.3 Disruptiveness

Disruptiveness evaluates how much a paper shifts its field’s trajectory using citation patterns. We employ the Disruption Index, which compares citations to the focal paper with its references to distinguish between papers that create new research directions versus those that consolidate existing research directions.

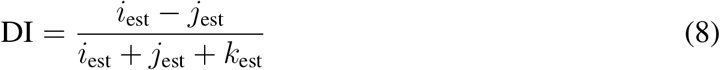

where *i*_est_ represents papers citing only the focal paper (indicating standalone impact), *j*_est_ represents papers citing only its references (reflecting influence of prior work), and *k*_est_ represents papers citing both (showing overlapping influence). These estimates are calculated from: (i) *A*, the focal paper’s citation count; (ii) *B*, the number of papers citing at least one reference; and (iii) *p*_*k*_, the proportion of sampled citing papers that also cite a reference. The estimates follow: *i*_est_ = (1 *− p*_*k*_) · *A, k*_est_ = *p*_*k*_ *· A*, and *j*_est_ = *B − k*_est_.

The index ranges from -1 to 1, where positive values indicate disruptive papers that garner citations independently thus overshadowing predecessors, negative values indicate consolidating papers that gathers citation alonside its predecessors, and zero represents neutral impact.

#### S6.4 Normalization

Novelty and disruptiveness measures are scaled quarterly across all universities to ensure temporal comparability and account for shifts in the research landscape that could affect absolute measure ranges:

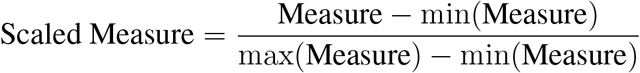

where min(Measure) and max(Measure) represent the minimum and maximum values across all universities in a given quarter. This normalization transforms raw measures to a [0, 1] range within each quarter. The pivot score requires no additional scaling as it is inherently normalized through cosine similarity.

### S7 Supplementary Figures

**Figure S1:**
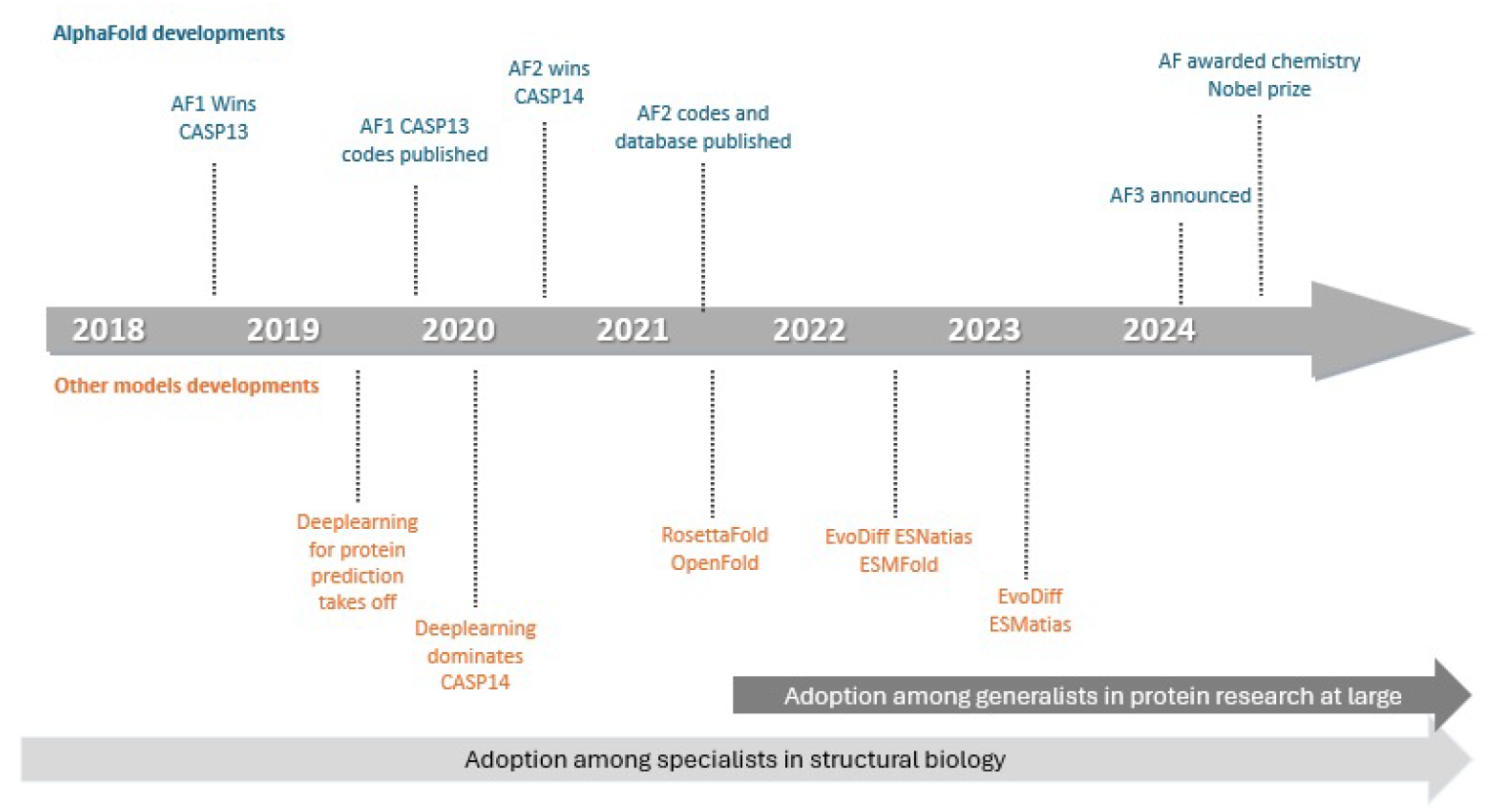
Timeline of AI development in structural biology. The development of AI systems in structural biology took place in two distinct phase. An early period, starting after the introduction of of AlphaFold 1 (AF1), where the adoption of AI likely requires substantial domain knowledge and ICT skills. Then a later phase with the introduction of the AlphaFold 2 (AF2) database where adoption becomes much easier and likely touch a much wider group. A third version, AlphaFold 3 (AF3) was recently announced 2024.

**Figure S2:**
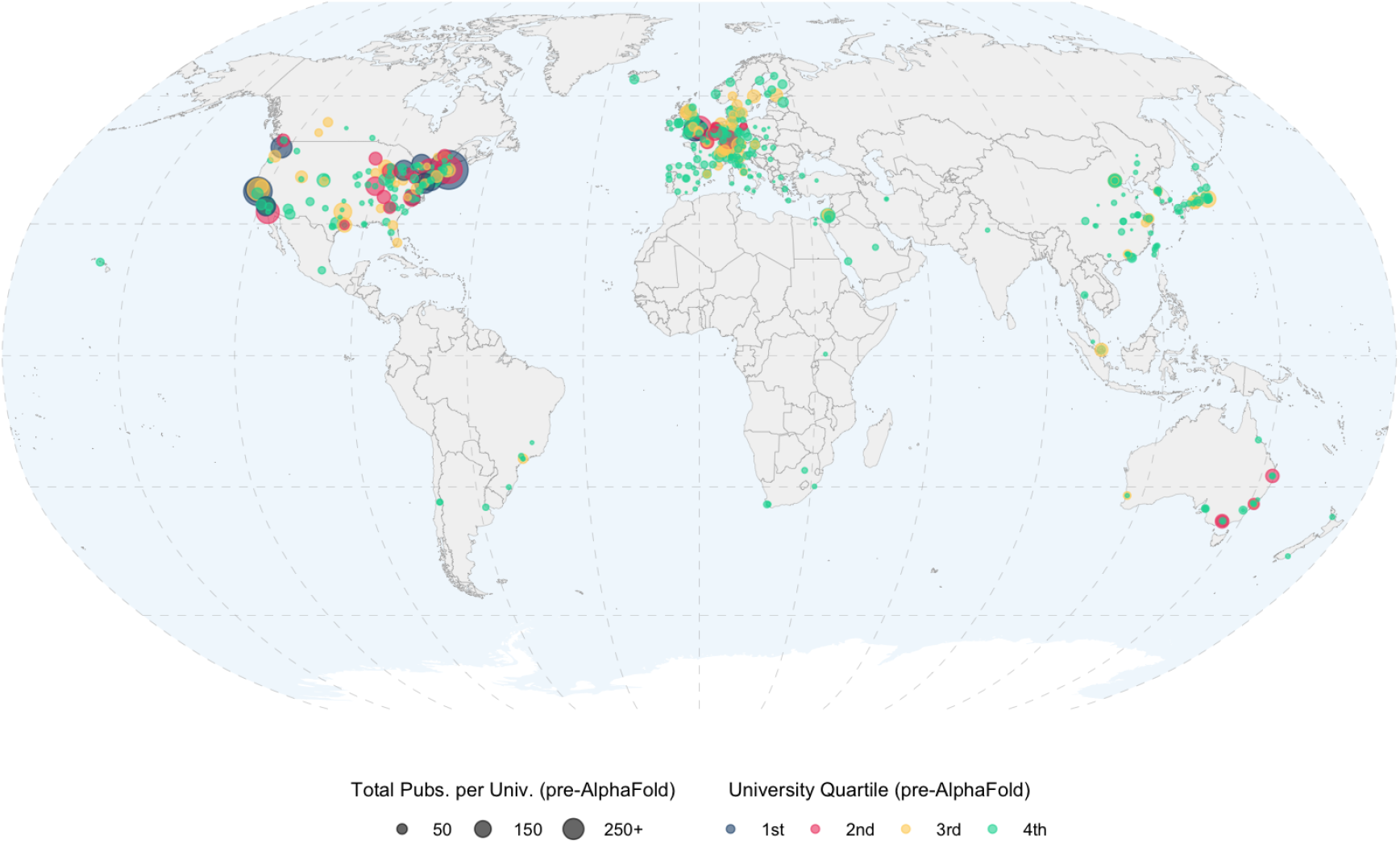
Geographical distribution of universities by quartile. Figure shows the location of the top 500 universities in our sample. Different colors represent different quartiles. Quartile allocation is based on 2013-2018 top journal market share. The diameter of each circle is based on the number of top journal publications a university recorded in the 2013-2018 period.

**Figure S3:**
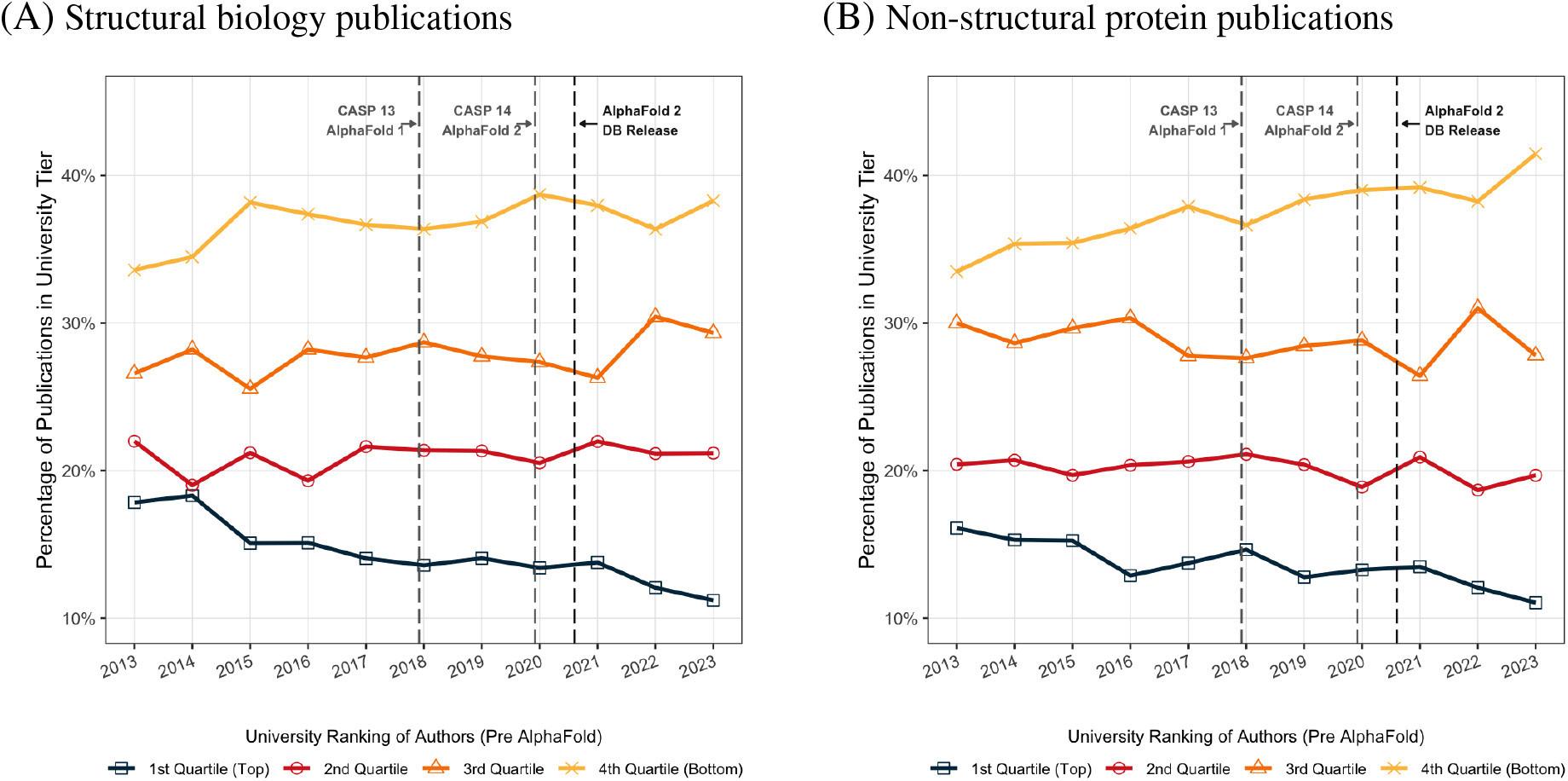
Publications in lower-tier journals by university quartiles. Figure illustrates the percentage of protein research publications in lower-tier journals with an SJR ∈ [1, 10) between 2013 and 2018, before the release of AlphaFold 1. The analysis is limited to authors affiliated with the top 500 universities, which are divided into four quartiles based on their aggregate share of publications in high-impact journals during the same period.

**Figure S4:**
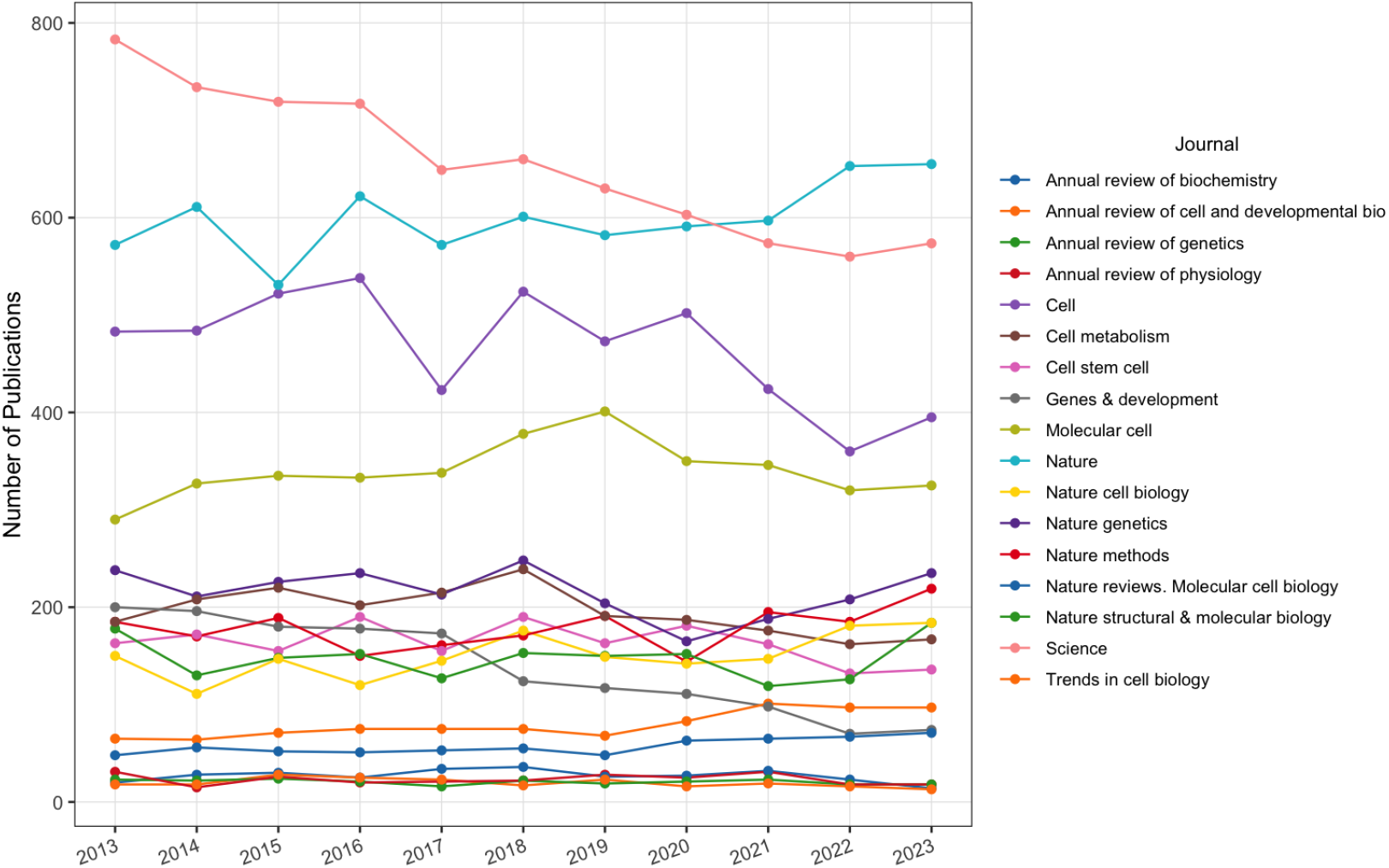
Distribution of top journal publications over time. Figure illustrates the annual number of research articles published in top-tier journals (SJR ≥10) between 2013 and 2023, covering three broad areas: structural biology, non-structural biology protein research, and non-protein research.

**Figure S5:**
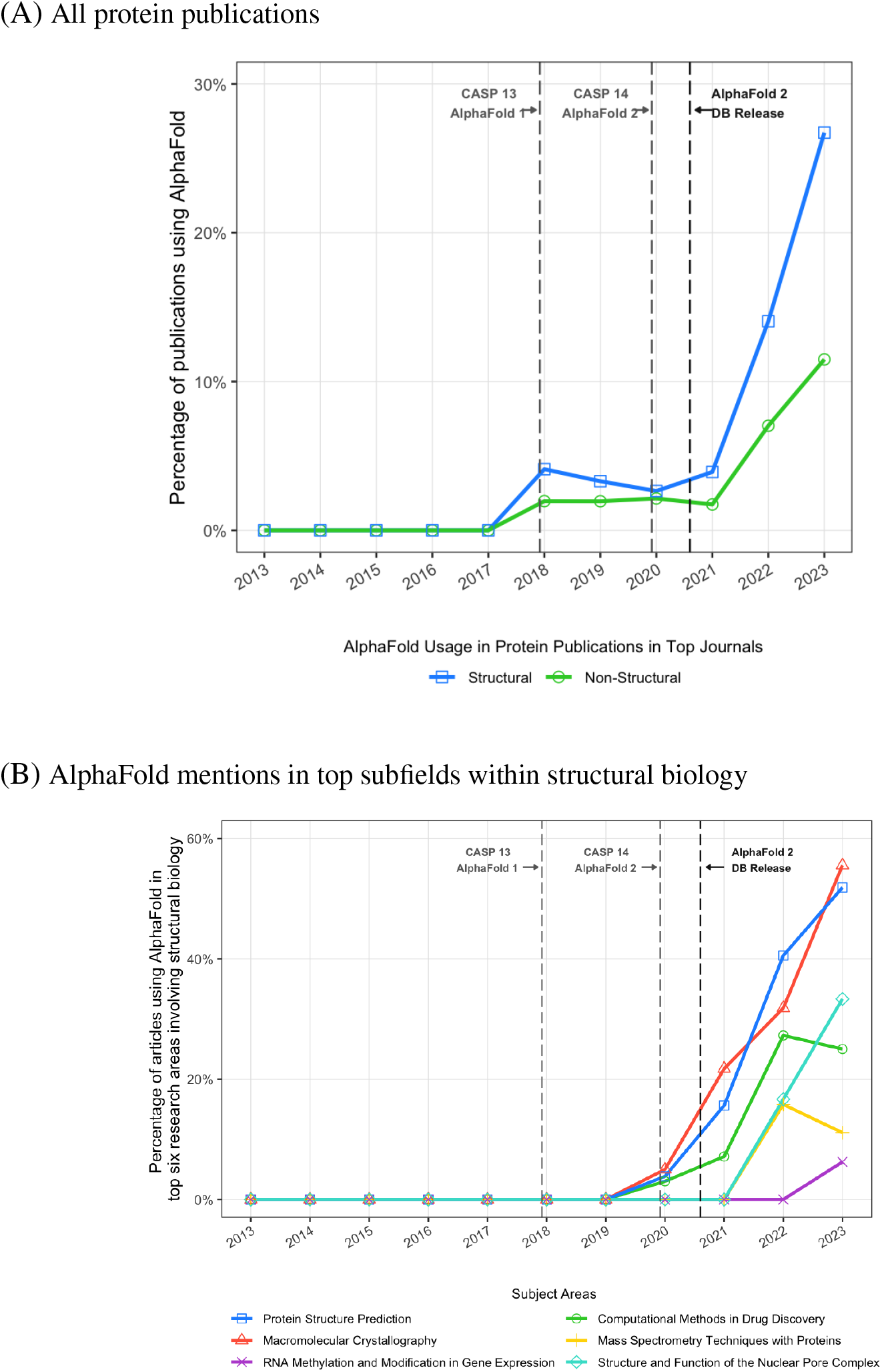
AlphaFold mentions in top protein publications. Figure shows details on AlphaFold adoption in top journals (SJR ≥10). Figure shows the share of protein research papers published that mention explicit use of AlphaFoldin their abstract or related AI tools, such as RoseTTAfold. Panel (A) shows this share for structural and non-structural protein research. Panel (B) breaks down the adoption rate by subfield of structural biology.

**Figure S6:**
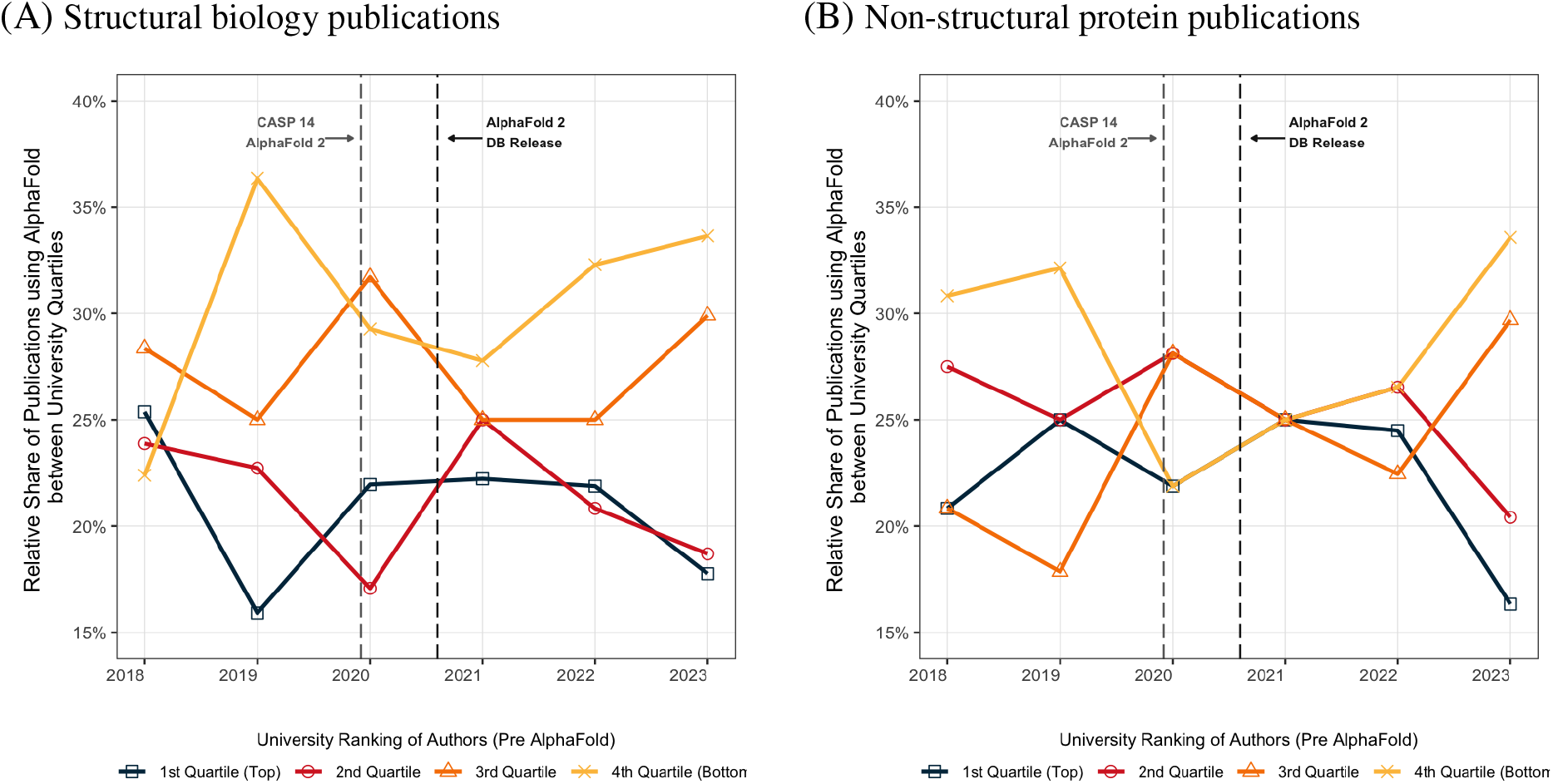
AlphaFold mentions in top protein publications by university quartile. Figure shows the relative share of AlphaFold usage among university quartiles in structural biology (Panel A) and non-structural protein publications (Panel B).

**Figure S7:**
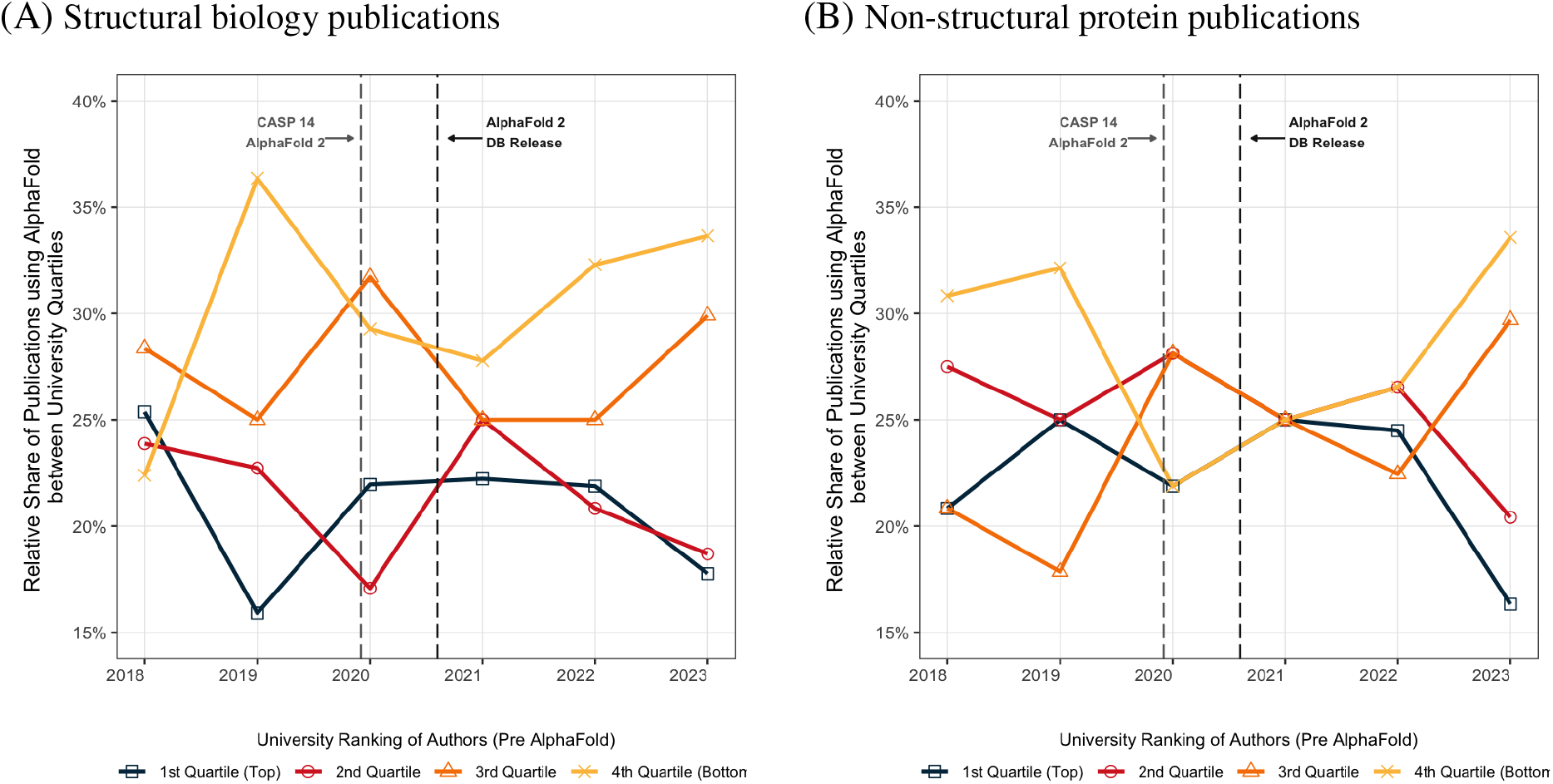
AlphaFold mentions in top protein publications by university quartile. Figure shows the relative share of AlphaFold usage among university quartiles in structural biology (Panel A) and non-structural protein publications (Panel B).

**Figure S8:**
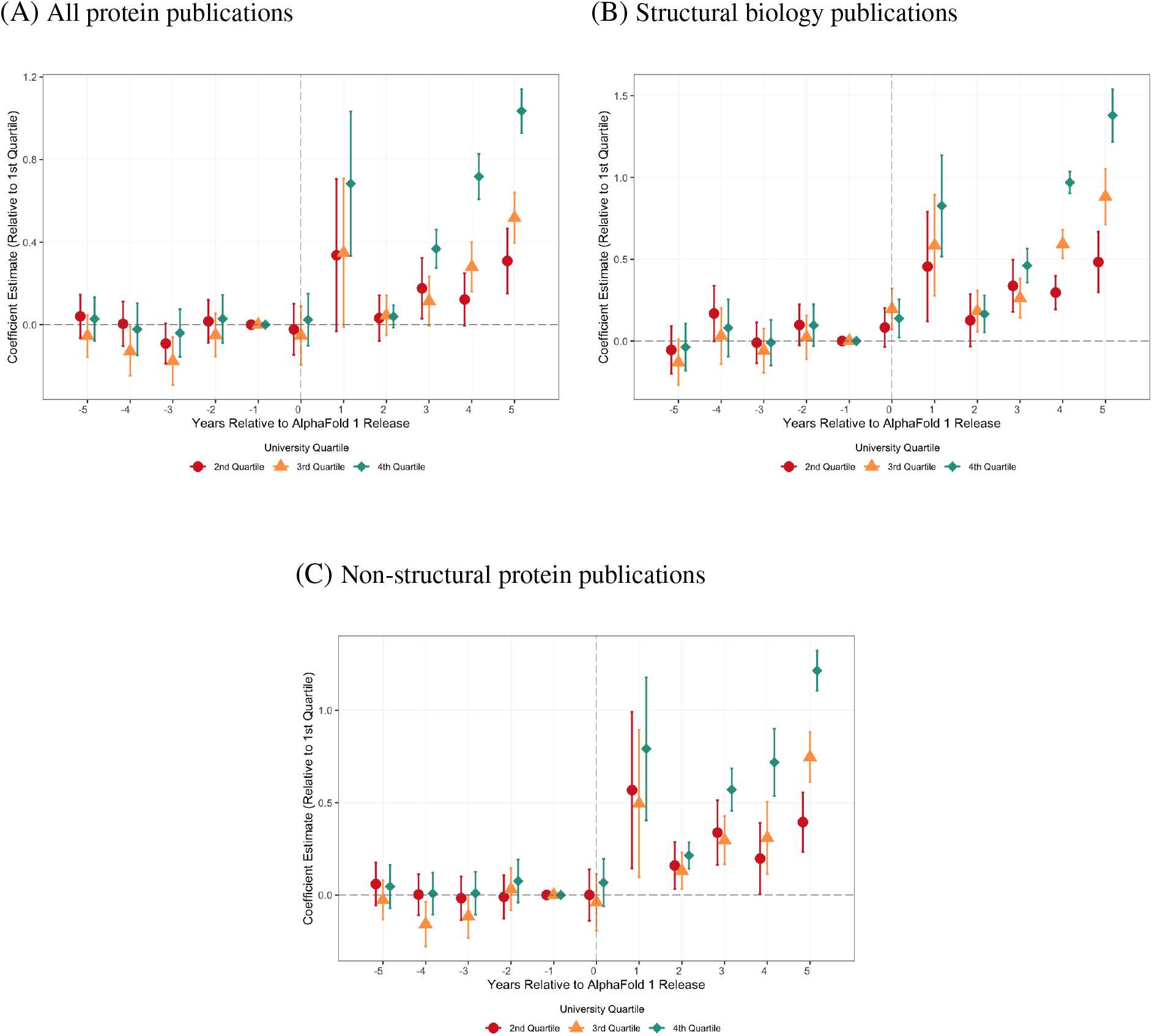
Protein publications by universities in top journals around AlphaFold : dynamic estimates. Figure illustrates yearly estimates obtained from regressing the number of protein publications in top journal on university quartile dummies, and the set of controls from our main specification. The plot shows point estimates with 95% confidence intervals. The dotted line represents the release date of AlphaFold. The analysis includes all protein publications (Panel A), structural biology publications (Panel B), and non-structural protein publications (Panel C). Universities are divided into four quartiles based on their aggregate share of publications in high-impact journals during 2013-2018.

**Figure S9:**
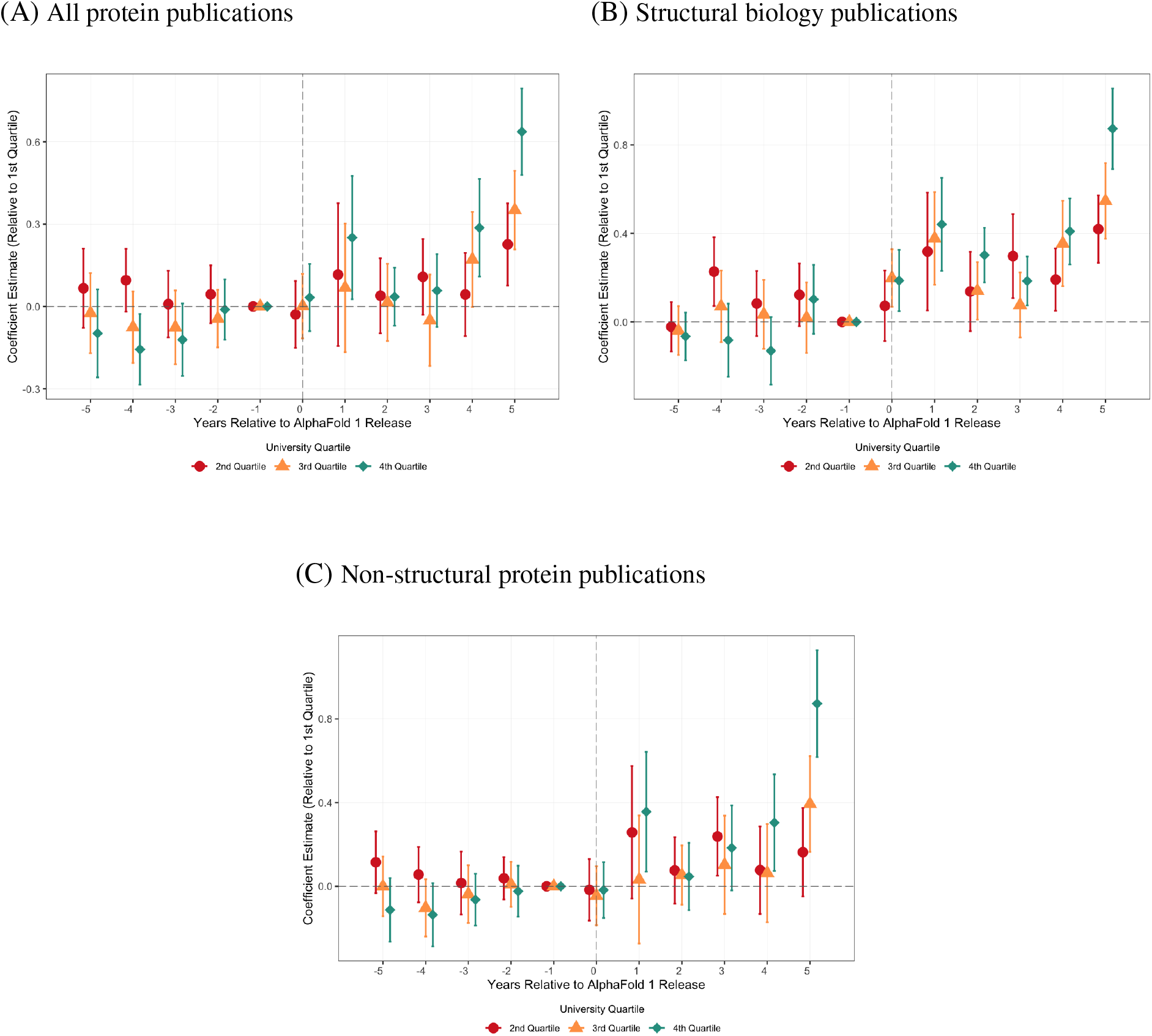
Market shares of universities in top journal protein publications around AlphaFold : dynamic estimates. Figure illustrates yearly estimates obtained from regressing top journal share on university quartile dummies, and the set of controls from our main specification. The plot shows point estimates with 95% confidence intervals. The dotted line represents the release date of AlphaFold. The analysis includes all protein publications (Panel A), structural biology publications (Panel B), and non-structural protein publications (Panel C). Universities are divided into four quartiles based on their aggregate share of publications in high-impact journals during 2013-2018.

### S8 Supplementary Tables

**Table S1:** Other AI tools developed for protein structure prediction. Table lists some of the most prominent AI tools that were developed following AlphaFold ‘s success. Though most of them are based on the same core transformer models like AlphaFold, many of them focus on different tasks and/or use different training data.

| AI Product | Release Date | Open-Source | Creator(s) | Key Feature |
| --- | --- | --- | --- | --- |
| RoseTTAFold | July 2021 | Yes | University of Washington | Predicts Protein-nucleic acid complexes |
| EvoDiff | September 2022 | Yes | Microsoft | Applies diffusion models to get protein design from evolutionary information |
| EsmFold | November 2022 | No | Meta AI | Uses amino acid sequence only for faster predictions. ESM atlas is a database with 617 million predicted protein structures |
| ProGen | January 2023 | No | Salesforce AI Research | Synthesizes new protein sequences with desired properties from natural language inputs |
| BaseFold | February 2023 | No | Basecamp Research | Uses proprietary protein samples in training data |
| RFDiffusion | July 2023 | Yes | University of Washington | Applies diffusion models to focus on protein design |

**Table S2:** Top journals in protein research. Table lists all journals with an SJR above ten which are used to constitute the sample of publication in the analysis throughout.

| Journal Name | ISSN | Mean SJR<br>(2013–18) | Mean Impact Factor<br>(2013–18) | Publisher |
| --- | --- | --- | --- | --- |
| Nature reviews. Molecular cell biology | 1471-0072 | 28.16 | 39.74 | Nature Portfolio |
| Cell | 0092-8674 | 27.21 | 32.02 | Cell Press |
| Nature genetics | 1061-4036 | 23.01 | 28.02 | Nature Portfolio |
| Annual rev. of biochemistry | 1545-4509 | 22.12 | 28.44 | Annual Reviews |
| Nature | 1476-4687 | 18.46 | 41.24 | Nature Portfolio |
| Nature methods | 1548-7091 | 18.14 | 27.13 | Nature Portfolio |
| Annual review of cell and dev. biology | 1530-8995 | 14.97 | 14.94 | Annual Reviews |
| Nature cell biology | 1476-4679 | 14.63 | 18.37 | Nature Portfolio |
| Cell stem cell | 1934-5909 | 13.70 | 22.49 | Elsevier BV |
| Molecular cell | 1097-4164 | 13.47 | 14.24 | Elsevier BV |
| Science | 1095-9203 | 13.08 | 39.10 | AAAS |
| Annual review of genetics | 1545-2948 | 12.95 | 11.98 | Annual Reviews |
| Cell metabolism | 1550-4131 | 11.21 | 18.79 | Cell Press |
| Genes & development | 1549-5477 | 11.06 | 10.84 | Cold Spring Harbor |
| Nature structural & molecular biology | 1545-9993 | 10.82 | 12.72 | Nature Portfolio |
| Trends in cell biology | 0962-8924 | 10.41 | 14.39 | Elsevier BV |
| Annual review of physiology | 0066-4278 | 10.04 | 17.22 | Annual Reviews |

**Table S3:** Descriptive statistics: top protein publications and university characteristics before AlphaFold. Table presents summary statistics at the university-year level for the period 2013–18, representing the years before the release of AlphaFold 1. The sample is restricted to protein-related publications in top journals with an SJR score of 10 or higher, authored by researchers affiliated with the top 500 universities prior to AlphaFold’s release. Each cell reports the mean of the respective variable, measured annually during the pre-AlphaFold period, with universities grouped into quartiles.

| Variable (*Pre AlphaFold estimates*) | 1<sup>st</sup> (reference group) | 2<sup>nd</sup> | 3<sup>rd</sup> | 4<sup>th</sup> |
| --- | --- | --- | --- | --- |
| ***Structural biology publications*** |  |  |  |  |
| Number of Publications per Year per University | 8.58 | 2.39 | 1.75 | 0.80 |
| Market Share (% publications) | 0.37 | 0.10 | 0.08 | 0.03 |
| Market Share (% of authors on publication) | 0.39 | 0.08 | 0.06 | 0.02 |
| ***Non-structural protein publications*** |  |  |  |  |
| Number of Publications per Year per University | 7.88 | 2.58 | 1.59 | 0.88 |
| Market Share (% publications) | 0.36 | 0.12 | 0.07 | 0.04 |
| Market Share (% of authors on publication) | 0.37 | 0.10 | 0.05 | 0.03 |
| ***University characteristics*** |  |  |  |  |
| Faculty Size | 614 | 308 | 267 | 178 |
| Share Structural Biologists (%) | 5.98 | 4.90 | 5.19 | 3.93 |
| Share Female (%) | 31.04 | 32.96 | 31.46 | 34.23 |
| Share White (%) | 28.91 | 26.11 | 23.62 | 22.61 |
| Share Asian (%) | 51.09 | 50.15 | 55.80 | 54.69 |
| Share Black (%) | 12.68 | 12.92 | 8.96 | 9.43 |
| Share affiliated in US (%) | 45.01 | 22.60 | 16.93 | 21.87 |
| Share affiliated in Europe (%) | 38.04 | 54.28 | 48.36 | 44.56 |
| Share affiliated in Asia (%) | 20.68 | 19.77 | 33.26 | 26.14 |
| Share affiliated with Govt (%) | 5.76 | 9.58 | 6.59 | 7.18 |
| Share affiliated with Research Lab (%) | 14.26 | 16.73 | 12.06 | 13.31 |
| Share affiliated with Industry (%) | 3.22 | 2.70 | 2.70 | 1.99 |
| Share affiliated with Medical Inst (%) | 16.97 | 16.92 | 16.46 | 14.46 |
| Share HPC Access (%) | 50.90 | 32.26 | 31.00 | 9.07 |
| HPC power (petaFlop/s) | 351.11 | 205.55 | 231.27 | 54.18 |
| Number of Universities | 12 | 36 | 88 | 364 |

**Table S4:** List of universities in the top quartile. These 12 universities together make up approximately 25% of top journal publications before AlphaFold (year 2013-2018)

| Name | Country |
| --- | --- |
| Harvard University | USA |
| Johns Hopkins University | USA |
| University of California, San Francisco | USA |
| University of Toronto | Canada |
| University of Pennsylvania | USA |
| University of Washington | USA |
| University College London | UK |
| Stanford University | USA |
| Duke University | USA |
| University of Michigan–Ann Arbor | USA |
| University of California, Los Angeles | USA |
| University of Oxford | UK |

**Table S5:** Descriptive statistics: top non-protein publications and university characteristics before AlphaFold. Table presents summary statistics at the university-year level for the period 2013–18, representing the years before the release of AlphaFold 1. The sample is restricted to non-protein publications in top journals with an SJR score of 10 or higher, authored by researchers affiliated with the top 500 universities prior to AlphaFold’s release. Each cell reports the mean and standard deviation (in brackets) of the respective variable, measured annually during the pre-AlphaFold period, with universities grouped into quartiles. ***, **, and * denote statistical significance in the difference in means between a variable in the given quartile and the first quartile (reference group) at the 1%, 5%, and 10% levels, respectively.

| Variable (*Pre AlphaFold estimates*) | University Quartile: mean (SD) |  |  |  |
| --- | --- | --- | --- | --- |
|  | 1<sup>st</sup> | 2<sup>nd</sup> | 3<sup>rd</sup> | 4<sup>th</sup> |
| Number of publications per year per university | 3.19(12.13) | 3.55(11) | 3.7(7.45)* | 2.89(4.86) |
| Market Share (% publications) | 0.06(0.25) | 0.07(0.22) | 0.07(0.15)* | 0.06(0.1) |
| Market share (% of authors on publication) | 0.06(0.28) | 0.08(0.27) | 0.07(0.16)* | 0.05(0.1) |
| Number of Universities | 12 | 36 | 88 | 364 |

**Table S6:** Structural biology top publications around AlphaFold: robustness checks. *PostAF1* is a binary variable set to one for papers published after the release of AlphaFold V1 in December 2018. The sample is limited to structural biology papers published in top journals with an SJR score of 10 or higher and to authors affiliated with the top 500 universities prior to the release of AlphaFold 1. Universities are ranked based on their average TNCS in the period preceding the release of AlphaFold 1. The highest ranked universities placed in the top quartile (*University Tier = 1st Quartile*) are used as the reference group. Standard errors are clustered by university, and reported in parentheses. ***, **, and * denote significance at the 1%, 5%, and 10% level, respectively.

| Dependent Variable: | Log(1 + No. of Publications) |  |  |  |  |
| --- | --- | --- | --- | --- | --- |
| Model: | (1) | (2) | (3) | (4) | (5) |
| University Tier = 2nd Quartile × PostAF1 | 0.2444***<br>(0.0172) | 0.2443***<br>(0.0173) | 0.2351***<br>(0.0145) | 0.2418***<br>(0.0197) | 0.2309***<br>(0.0166) |
| University Tier = 3rd Quartile × PostAF1 | 0.3555***<br>(0.0211) | 0.3557***<br>(0.0210) | 0.3764***<br>(0.0262) | 0.3425***<br>(0.0272) | 0.3605***<br>(0.0322) |
| University Tier = 4th Quartile × PostAF1 | 0.5849***<br>(0.0321) | 0.5845***<br>(0.0320) | 0.5598***<br>(0.0334) | 0.5678***<br>(0.0379) | 0.5392***<br>(0.0412) |
| Controls: Share Structural Biologists |  | ✓ |  |  | ✓ |
| Controls: HPC Power |  |  | ✓ |  | ✓ |
| Controls: Author Characteristics |  |  |  | ✓ | ✓ |
| Year FE | ✓ | ✓ | ✓ | ✓ | ✓ |
| University FE | ✓ | ✓ | ✓ | ✓ | ✓ |
| Observations | 5,500 | 5,500 | 5,500 | 5,500 | 5,500 |
| Adjusted R<sup>2</sup> | 0.780 | 0.781 | 0.805 | 0.785 | 0.809 |

(B) Market share (% publications, all authors)
| Dependent Variable: | Market Share (% Publications, All Authors) |  |  |  |  |
| --- | --- | --- | --- | --- | --- |
| Model: | (1) | (2) | (3) | (4) | (5) |
| University Tier = 2nd Quartile × PostAF1 | 0.1387***<br>(0.0019) | 0.1394***<br>(0.0018) | 0.1406***<br>(0.0014) | 0.1322***<br>(0.0049) | 0.1342***<br>(0.0037) |
| University Tier = 3rd Quartile × PostAF1 | 0.1433***<br>(0.0028) | 0.1445***<br>(0.0025) | 0.1738***<br>(0.0219) | 0.1404***<br>(0.0059) | 0.1644***<br>(0.0216) |
| University Tier = 4th Quartile × PostAF1 | 0.2827***<br>(0.0043) | 0.2813***<br>(0.0050) | 0.2837***<br>(0.0232) | 0.2584***<br>(0.0105) | 0.2497***<br>(0.0271) |
| Controls: Share Structural Biologists |  | ✓ |  |  | ✓ |
| Controls: HPC Power |  |  | ✓ |  | ✓ |
| Controls: Author Characteristics |  |  |  | ✓ | ✓ |
| Year FE | ✓ | ✓ | ✓ | ✓ | ✓ |
| University FE | ✓ | ✓ | ✓ | ✓ | ✓ |
| Observations | 5,500 | 5,500 | 5,500 | 5,500 | 5,500 |
| Pseudo R<sup>2</sup> | 0.904 | 0.904 | 0.906 | 0.917 | 0.921 |

(C) Market share (% publications, main authors)
| Dependent Variable: | Market Share (% Publications, Main Authors) |  |  |  |  |
| --- | --- | --- | --- | --- | --- |
| Model: | (1) | (2) | (3) | (4) | (5) |
| University Tier = 2nd Quartile × PostAF1 | 0.1482***<br>(0.0020) | 0.1480***<br>(0.0024) | 0.1291***<br>(0.0029) | 0.1683***<br>(0.0036) | 0.1480***<br>(0.0043) |
| University Tier = 3rd Quartile × PostAF1 | 0.1409***<br>(0.0032) | 0.1405***<br>(0.0036) | 0.1543***<br>(0.0160) | 0.1578***<br>(0.0063) | 0.1623***<br>(0.0145) |
| University Tier = 4th Quartile × PostAF1 | 0.2700***<br>(0.0044) | 0.2664***<br>(0.0058) | 0.2688***<br>(0.0209) | 0.2560***<br>(0.0111) | 0.2420***<br>(0.0242) |
| Controls: Share Structural Biologists |  | ✓ |  |  | ✓ |
| Controls: HPC Power |  |  | ✓ |  | ✓ |
| Controls: Author Characteristics |  |  |  | ✓ | ✓ |
| Year FE | ✓ | ✓ | ✓ | ✓ | ✓ |
| University FE | ✓ | ✓ | ✓ | ✓ | ✓ |
| Observations | 5,500 | 5,500 | 5,500 | 5,500 | 5,500 |
| Pseudo R<sup>2</sup> | 0.899 | 0.901 | 0.903 | 0.915 | 0.919 |

**Table S7:** Non-structural protein top publications around AlphaFold: robustness checks. *PostAF1* is a binary variable set to one for papers published after the release of AlphaFold 1 in December 2018. The sample is limited to non-structural biology protein papers published in top journals with an SJR score of 10 or higher and to authors affiliated with the top 500 universities prior to the release of AlphaFold 1. Universities are ranked based on their average TNCS in the period preceding the release of AlphaFold 1. The highest ranked universities placed in the top quartile (*University Tier = 1st Quartile*) are used as the reference group. Standard errors are clustered by university, and reported in parentheses. ***, **, and * denote significance at the 1%, 5%, and 10% level, respectively.

| Dependent Variable: | Log(1 + No. of Publications) |  |  |  |  |
| --- | --- | --- | --- | --- | --- |
| Model: | (1) | (2) | (3) | (4) | (5) |
| University Tier = 2nd Quartile × PostAF1 | 0.2912***<br>(0.0201) | 0.2909***<br>(0.0201) | 0.2931***<br>(0.0194) | 0.2757***<br>(0.0281) | 0.2767***<br>(0.0276) |
| University Tier = 3rd Quartile × PostAF1 | 0.4052***<br>(0.0185) | 0.4043***<br>(0.0185) | 0.4183***<br>(0.0215) | 0.3913***<br>(0.0240) | 0.4018***<br>(0.0289) |
| University Tier = 4th Quartile × PostAF1 | 0.5963***<br>(0.0279) | 0.5957***<br>(0.0279) | 0.5572***<br>(0.0259) | 0.5766***<br>(0.0347) | 0.5361***<br>(0.0340) |
| Controls: Share Structural Biologists |  | ✓ |  |  | ✓ |
| Controls: HPC Power |  |  | ✓ |  | ✓ |
| Controls: Author Characteristics |  |  |  | ✓ | ✓ |
| Year FE | ✓ | ✓ | ✓ | ✓ | ✓ |
| University FE | ✓ | ✓ | ✓ | ✓ | ✓ |
| Observations | 5,500 | 5,500 | 5,500 | 5,500 | 5,500 |
| Adjusted R<sup>2</sup> | 0.760 | 0.761 | 0.787 | 0.765 | 0.791 |

(B) Market share (% publications, all authors)
| Dependent Variable: | Market Share (% Publications, all authors) |  |  |  |  |
| --- | --- | --- | --- | --- | --- |
| Model: | (1) | (2) | (3) | (4) | (5) |
| University Tier = 2nd Quartile × PostAF1 | 0.0876***<br>(0.0028) | 0.0869***<br>(0.0023) | 0.0943***<br>(0.0061) | 0.0840***<br>(0.0050) | 0.0904***<br>(0.0083) |
| University Tier = 3rd Quartile × PostAF1 | 0.1482***<br>(0.0030) | 0.1488***<br>(0.0032) | 0.1834***<br>(0.0195) | 0.1392***<br>(0.0084) | 0.1715***<br>(0.0280) |
| University Tier = 4th Quartile × PostAF1 | 0.2505***<br>(0.0044) | 0.2501***<br>(0.0044) | 0.2628***<br>(0.0143) | 0.2387***<br>(0.0074) | 0.2470***<br>(0.0227) |
| Controls: Share Structural Biologists |  | ✓ |  |  | ✓ |
| Controls: HPC Power |  |  | ✓ |  | ✓ |
| Controls: Author Characteristics |  |  |  | ✓ | ✓ |
| Year FE | ✓ | ✓ | ✓ | ✓ | ✓ |
| University FE | ✓ | ✓ | ✓ | ✓ | ✓ |
| Observations | 5,500 | 5,500 | 5,500 | 5,500 | 5,500 |
| Pseudo R<sup>2</sup> | 0.884 | 0.883 | 0.888 | 0.899 | 0.902 |

(C) Market share (% publications, main authors)
| Dependent Variable: | Market Share (% Publications, main authors) |  |  |  |  |
| --- | --- | --- | --- | --- | --- |
| Model: | (1) | (2) | (3) | (4) | (5) |
| University Tier = 2nd Quartile × PostAF1 | 0.1246***<br>(0.0024) | 0.1242***<br>(0.0017) | 0.1288***<br>(0.0052) | 0.1142***<br>(0.0070) | 0.1164***<br>(0.0096) |
| University Tier = 3rd Quartile × PostAF1 | 0.1532***<br>(0.0024) | 0.1529***<br>(0.0019) | 0.1893***<br>(0.0195) | 0.1398***<br>(0.0131) | 0.1707**<br>(0.0312) |
| University Tier = 4th Quartile × PostAF1 | 0.2840***<br>(0.0041) | 0.2832***<br>(0.0038) | 0.3096***<br>(0.0185) | 0.2767***<br>(0.0117) | 0.2962***<br>(0.0300) |
| Controls: Share Structural Biologists |  | ✓ |  |  | ✓ |
| Controls: HPC Power |  |  | ✓ |  | ✓ |
| Controls: Author Characteristics |  |  |  | ✓ | ✓ |
| Year FE | ✓ | ✓ | ✓ | ✓ | ✓ |
| University FE | ✓ | ✓ | ✓ | ✓ | ✓ |
| Observations | 5,500 | 5,500 | 5,500 | 5,500 | 5,500 |
| Pseudo R<sup>2</sup> | 0.881 | 0.880 | 0.885 | 0.897 | 0.901 |

**Table S8:** Research patterns in top journal publications around AlphaFold. Table presents regression estimates examining qualitative changes in the research produced following AlphaFold 1 release. Dependent variables are pivot scores, novelty measures, and disruptiveness scores for publications in top journals (SJR ≥ 10). *PostAF1* is a binary indicator for papers published after December 2018. Sample includes structural biology publications (columns 1-3) and non-structural protein publications (columns 4-6) by authors affiliated with top 500 universities. University quartiles based on pre-AlphaFold publication shares in high-impact journals (2013-2018), with 1st quartile as reference group. All measures scaled quarterly within [0, 1] range for temporal comparability. Standard errors clustered by university in parentheses. ***, **, * denote significance at 1%, 5%, 10% levels.

| Dependent Variables:<br>Model: | Structural Biology Pubs. |  |  | Non-Structural Protein Pubs. |  |  |
| --- | --- | --- | --- | --- | --- | --- |
|  | Pivot Score | Novelty | Disruptiveness | Pivot Score | Novelty | Disruptiveness |
|  | (1) | (2) | (3) | (4) | (5) | (6) |
| University Tier = 2nd Quartile × PostAF1 | 0.0104**<br>(0.0025) | 0.0196***<br>(0.0020) | 0.0117***<br>(0.0003) | 0.0345***<br>(0.0013) | 0.0026<br>(0.0015) | -0.0010<br>(0.0012) |
| University Tier = 3rd Quartile × PostAF1 | 0.0166***<br>(0.0026) | 0.0152***<br>(0.0020) | 0.0046**<br>(0.0010) | 0.0028<br>(0.0025) | 0.0112***<br>(0.0011) | -0.0056*<br>(0.0022) |
| University Tier = 4th Quartile × PostAF1 | 0.0263***<br>(0.0021) | 0.0154**<br>(0.0032) | 0.0126***<br>(0.0015) | 0.0326***<br>(0.0034) | 0.0268***<br>(0.0018) | 0.0065*<br>(0.0026) |
| Year FE | ✓ | ✓ | ✓ | ✓ | ✓ | ✓ |
| University FE | ✓ | ✓ | ✓ | ✓ | ✓ | ✓ |
| Observations | 5,500 | 5,500 | 5,500 | 5,500 | 5,500 | 5,500 |
| R<sup>2</sup> | 0.03 | 0.18 | 0.07 | 0.02 | 0.16 | 0.06 |

**Table S9:** Collaborations with top-quartile universities in top journal publications around AlphaFold. Table presents regression estimates examining changes in collaboration patterns following AlphaFold 1 release. Dependent variable is the share of top-journal publications produced by university *u* in year *t* that were co-authored with at least one author from a top-quartile university. For each sample, we run two specifications: One were we only include the usual double interaction term between *PostAF1* and *University Tier* (column 1,3,5) and one where we include the triple interaction term between *PostAF1, University Tier*, and the share of structural biologists at the university (column 2,4,6) in year t. University quartiles based on pre-AlphaFold publication shares in high-impact journals (2013-2018), with 1st quartile as reference group. *PostAF1* is a binary indicator for papers published after December 2018 Quartile 1 is the reference group and the controls are the same as in Table S3.

| Dependent Variable: | Collaboration 1st Quartile (%) |  |  |  |  |  |
| --- | --- | --- | --- | --- | --- | --- |
|  | All Protein Pubs. |  | Structural Biology Pubs. |  | Non-Structural Protein Pubs. |  |
| Model: | (1) | (2) | (3) | (4) | (5) | (6) |
| University Tier = 2nd Quartile × PostAF1 | -0.0124***<br>(0.0004) | 0.0738**<br>(0.0199) | -0.0491***<br>(0.0021) | 0.1535***<br>(0.0210) | -0.0337**<br>(0.0081) | 0.1704***<br>(0.0163) |
| University Tier = 3rd Quartile × PostAF1 | 0.0006<br>(0.0091) | 0.1093***<br>(0.0175) | -0.0657**<br>(0.0118) | 0.1678***<br>(0.0146) | -0.0506**<br>(0.0151) | 0.1544***<br>(0.0166) |
| University Tier = 4th Quartile × PostAF1 | -0.0566**<br>(0.0124) | 0.0377*<br>(0.0131) | -0.1295***<br>(0.0146) | 0.0937***<br>(0.0102) | -0.1162***<br>(0.0174) | 0.0665**<br>(0.0158) |
| University Tier = 2nd Quartile × PostAF1 × Structural Biologists (%) | | -1.240**<br>(0.2747) | | -2.750***<br>(0.3011) | | -2.788***<br>(0.2796) |
| University Tier = 3rd Quartile × PostAF1 × Structural Biologists (%) | | -1.432**<br>(0.3628) | | -3.030***<br>(0.3633) | | -2.815***<br>(0.3428) |
| University Tier = 4th Quartile × PostAF1 × Structural Biologists (%) | | -1.079**<br>(0.3363) | | -2.745***<br>(0.3140) | | -2.377***<br>(0.3096) |
| Controls | ✓ | ✓ | ✓ | ✓ | ✓ | ✓ |
| Year FE | ✓ | ✓ | ✓ | ✓ | ✓ | ✓ |
| University FE | ✓ | ✓ | ✓ | ✓ | ✓ | ✓ |
| Observations | 5,500 | 5,500 | 5,500 | 5,500 | 5,500 | 5,500 |
| Adjusted R<sup>2</sup> | 0.19 | 0.20 | 0.29 | 0.29 | 0.30 | 0.30 |

**Table S10:** Structural biology publications in top journals around AlphaFold: robustness to controlling for research patterns. Table presents estimates that investigate the mediating effect of collaboration and research patterns on the relationship between AI and publication outcomes for structural biology publications in top journals (SJR ≥10). Panel A presents results with publication volume as the dependent variable. Panel B presents results with market share as the dependent variable. The computation of novelty, disruptiveness and pivot scores is documented in Section S6. Collaboration is computed as the share of top journal publications produced by university *u* in year *t* that were co-authored with at least one author from a top-quartile university.

| Dependent Variable: | Log(1 + No. of Publications) |  |  |  |  |  |
| --- | --- | --- | --- | --- | --- | --- |
| Model: | (1) | (2) | (3) | (4) | (5) | (6) |
| University Tier = 2nd Quartile × PostAF1 | 0.2475***<br>(0.0201) | 0.2473***<br>(0.0205) | 0.2490***<br>(0.0201) | 0.2489***<br>(0.0207) | 0.2484***<br>(0.0224) | 0.2474***<br>(0.0211) |
| University Tier = 3rd Quartile × PostAF1 | 0.3790***<br>(0.0424) | 0.3789***<br>(0.0430) | 0.3793***<br>(0.0423) | 0.3790***<br>(0.0431) | 0.3944***<br>(0.0479) | 0.3759***<br>(0.0441) |
| University Tier = 4th Quartile × PostAF1 | 0.6369***<br>(0.0754) | 0.6368***<br>(0.0761) | 0.6386***<br>(0.0754) | 0.6383***<br>(0.0762) | 0.6291***<br>(0.0882) | 0.6341***<br>(0.0777) |
| *Controls for Research Patterns* |  |  |  |  |  |  |
| Pivoting | ✓ |  |  | ✓ |  | ✓ |
| Novelty |  | ✓ |  | ✓ |  | ✓ |
| Disruptiveness |  |  | ✓ | ✓ |  | ✓ |
| Collaboration with 1st Quartile (%) | ✓ |  |  |  | ✓ | ✓ |
| Controls | ✓ | ✓ | ✓ | ✓ | ✓ | ✓ |
| Year FE | ✓ | ✓ | ✓ | ✓ | ✓ | ✓ |
| University FE | ✓ | ✓ | ✓ | ✓ | ✓ | ✓ |
| Observations | 5,500 | 5,500 | 5,500 | 5,500 | 5,500 | 5,500 |
| Adjusted R<sup>2</sup> | 0.87 | 0.87 | 0.86 | 0.86 | 0.87 | 0.86 |

(B) Market Shares
| Dependent Variable: | Market Share (% Publications, All Authors) |  |  |  |  |  |
| --- | --- | --- | --- | --- | --- | --- |
| Model: | (1) | (2) | (3) | (4) | (5) | (6) |
| University Tier = 2nd Quartile × PostAF1 | 0.0626***<br>(0.0013) | 0.1404***<br>(0.0040) | 0.1412***<br>(0.0035) | 0.1407***<br>(0.0037) | 0.1457***<br>(0.0083) | 0.1369***<br>(0.0045) |
| University Tier = 3rd Quartile × PostAF1 | 0.0689***<br>(0.0046) | 0.1527***<br>(0.0055) | 0.1534***<br>(0.0056) | 0.1529***<br>(0.0061) | 0.1677**<br>(0.0293) | 0.1422***<br>(0.0084) |
| University Tier = 4th Quartile × PostAF1 | 0.1397***<br>(0.0106) | 0.3718***<br>(0.0154) | 0.3732***<br>(0.0170) | 0.3723***<br>(0.0165) | 0.2747**<br>(0.0569) | 0.3591***<br>(0.0186) |
| *Controls for Research Patterns* |  |  |  |  |  |  |
| Pivoting | ✓ |  |  | ✓ |  | ✓ |
| Novelty |  | ✓ |  | ✓ |  | ✓ |
| Disruptiveness |  |  | ✓ | ✓ |  | ✓ |
| Collaboration with 1st Quartile (%) | ✓ |  |  |  | ✓ | ✓ |
| Controls | ✓ | ✓ | ✓ | ✓ | ✓ | ✓ |
| Year FE | ✓ | ✓ | ✓ | ✓ | ✓ | ✓ |
| University FE | ✓ | ✓ | ✓ | ✓ | ✓ | ✓ |
| Observations | 5,500 | 5,500 | 5,500 | 5,500 | 5,500 | 5,500 |
| Pseudo R<sup>2</sup> | 0.94 | 0.95 | 0.95 | 0.95 | 0.94 | 0.95 |

**Table S11:** Non-structural protein publications in top journals around AlphaFold: robustness to controlling for research patterns. Table presents estimates that investigate the mediating effect of collaboration and research patterns on the relationship between AI and publication outcomes for Non-structural protein publications in top journals (SJR ≥10). Panel A presents results with publication volume as the dependent variable. Panel B presents results with market share as the dependent variable. The computation of novelty, disruptiveness and pivot scores is documented in Section S6. Collaboration is computed as the share of top journal publications produced by university *u* in year *t* that were co-authored with at least one author from a top-quartile university.

| Dependent Variable: | Log(1 + No. of Publications) |  |  |  |  |  |
| --- | --- | --- | --- | --- | --- | --- |
| Model: | (1) | (2) | (3) | (4) | (5) | (6) |
| University Tier = 2nd Quartile × PostAF1 | 0.3355***<br>(0.0500) | 0.3363***<br>(0.0507) | 0.3362***<br>(0.0505) | 0.3355***<br>(0.0503) | 0.3703***<br>(0.0516) | 0.3312***<br>(0.0510) |
| University Tier = 3rd Quartile × PostAF1 | 0.5008***<br>(0.0449) | 0.5014***<br>(0.0457) | 0.5005***<br>(0.0450) | 0.5008***<br>(0.0457) | 0.5136***<br>(0.0477) | 0.4954***<br>(0.0467) |
| University Tier = 4th Quartile × PostAF1 | 0.6525***<br>(0.0797) | 0.6537***<br>(0.0812) | 0.6540***<br>(0.0800) | 0.6543***<br>(0.0811) | 0.6669***<br>(0.0856) | 0.6471***<br>(0.0825) |
| *Controls for Research Patterns* |  |  |  |  |  |  |
| Pivoting | ✓ |  |  | ✓ |  | ✓ |
| Novelty |  | ✓ |  | ✓ |  | ✓ |
| Disruptiveness |  |  | ✓ | ✓ |  | ✓ |
| Collaboration with 1st Quartile (%) | ✓ |  |  |  | ✓ | ✓ |
| Controls | ✓ | ✓ | ✓ | ✓ | ✓ | ✓ |
| Year FE | ✓ | ✓ | ✓ | ✓ | ✓ | ✓ |
| University FE | ✓ | ✓ | ✓ | ✓ | ✓ | ✓ |
| Observations | 5,500 | 5,500 | 5,500 | 5,500 | 5,500 | 5,500 |
| Adjusted R<sup>2</sup> | 0.84 | 0.84 | 0.84 | 0.85 | 0.85 | 0.85 |

(B) Market Shares
| Dependent Variable: | Market Share (% Publications, All Authors) |  |  |  |  |  |
| --- | --- | --- | --- | --- | --- | --- |
| Model: | (1) | (2) | (3) | (4) | (5) | (6) |
| University Tier = 2nd Quartile × PostAF1 | 0.0466***<br>(0.0032) | 0.1012***<br>(0.0046) | 0.1011***<br>(0.0049) | 0.1006***<br>(0.0073) | 0.1128***<br>(0.0167) | 0.0927***<br>(0.0077) |
| University Tier = 3rd Quartile × PostAF1 | 0.0665***<br>(0.0039) | 0.1545***<br>(0.0095) | 0.1544***<br>(0.0097) | 0.1542***<br>(0.0106) | 0.1643**<br>(0.0341) | 0.1374***<br>(0.0120) |
| University Tier = 4th Quartile × PostAF1 | 0.1232***<br>(0.0068) | 0.3417***<br>(0.0100) | 0.3421***<br>(0.0100) | 0.3423***<br>(0.0115) | 0.2489**<br>(0.0537) | 0.3222***<br>(0.0146) |
| *Controls for Research Patterns* |  |  |  |  |  |  |
| Pivoting | ✓ |  |  | ✓ |  | ✓ |
| Novelty |  | ✓ |  | ✓ |  | ✓ |
| Disruptiveness |  |  | ✓ | ✓ |  | ✓ |
| Collaboration with 1st Quartile (%) | ✓ |  |  |  | ✓ | ✓ |
| Controls | ✓ | ✓ | ✓ | ✓ | ✓ | ✓ |
| Year FE | ✓ | ✓ | ✓ | ✓ | ✓ | ✓ |
| University FE | ✓ | ✓ | ✓ | ✓ | ✓ | ✓ |
| Observations | 5,500 | 5,500 | 5,500 | 5,500 | 5,500 | 5,500 |
| Pseudo R<sup>2</sup> | 0.93 | 0.94 | 0.94 | 0.94 | 0.93 | 0.94 |

## Footnotes

1 See section S2.2 for a formal definition of 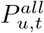 and 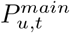

2 The research metrics *pivot behavior, novelty*, and *disruptiveness* are normalized to [0,1] for comparability. Methods used to compute these metrics are described in Section S6.

3 To motivate the analysis, we present some examples of how AlphaFold impacted various fields in protein research: AlphaFold has enabled the discovery of new 3D shapes (folds) in parts of viruses that bind to sugar molecules (glycan-binding domains), shifting research from studying rigid, crystal-like arrangements (paracrystalline) to more flexible, varied forms (polymorphic assembly in rotavirus studies [17]). In drug discovery, AlphaFold sped up the identification of mind-affecting drugs (psychotropic agonists) that target a brain protein involved in mood (serotonin receptor 1A), achieving a higher success rate (hit rate) than predictions based on similar known structures (traditional homology models) [18]. Additionally, AlphaFold supported predictions across all proteins in an organism (proteome-wide) for changes in DNA that alter protein building blocks (missense variant effects), improving understanding of their disease-causing potential (pathogenicity) and aiding gene-based medicine (clinical genomics) [19]. AlphaFold has also been instrumental to uncovering the structure of an immune signaling complex (IL-27), revealing new ways cells communicate during inflammation [20]. Access to AlphaFold also accelerated the discovery of a small molecule inhibitor for CDK20, a protein linked to cancer, by generating accurate structures for rapid screening [21].

4 For example, researchers at McMaster University (ranked 69 in our sample) and others used AlphaFold to study how proteins interact based on their genetic sequences, showing that the sequence is more important than the 3D structure for predicting these interactions in genomics [31]. This approach allowed them to incorporate uncommon ideas about protein behavior that were hard to test previously, boosting the originality of their findings, and helped better understand how enzymes repair DNA in bacteria. Similarly, researchers at the University of Alberta (ranked 84 in our sample) incorporated AlphaFold predictions to identify molecular markers for better medical diagnosis and treatment, enabling exploration of rare protein functions in bacterial genomes by adding notes to predicted structures in their system, which enhanced the novelty of their research by uncovering new biological insights [32]. Additionally, researchers at Universidade de Lisboa (ranked 315 in our sample) used AlphaFold to model a protein structure (CDS2), revealing potential spots for drugs to target in cancer-related pathways, and to design small binders for a receptor involved in cell processes for targeted breakdown [33]. Their approach introduced original concepts in combining structure with function for new therapeutic ideas. Lastly, researchers at University of Coimbra (ranked 372 in our sample) participated in a study that applied AlphaFold to predict a protein structure (TEX264), identifying parts involved in clearing damaged DNA, which brought fresh perspectives to DNA repair mechanisms and increased the work’s originality by linking structure to rare cellular cleanup processes [34].

5 While our discussion focuses on university market shares in top journal protein publications, a parallel analysis using raw annual publication volume as the dependent variable yields consistent results (see Tables S10(A) and S11(A)).

6 The coefficients from the logit model are log-odds ratios. The percentage change in the odds ratio is calculated as 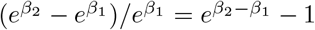, where *β*_1_ and *β*_2_ are the log-odds ratios before and after adding controls.

7 AF2 scored above 90 for around two-thirds of the proteins. This score represents the degree to which a predicted structure (such as using a computational tool like AlphaFold) is similar to the experimentally determined structure, with 100 denoting a complete match between the two. AF2 achieved a world record-breaking overall score of 92.4, representing exceptional precision in protein structure prediction See https://www.guinnessworldrecords.com/world-records/642132-highest-score-at-the-casp-competition.

8 Microsoft Academic Graph is less comprehensive than Google Scholar but more comprehensive than other providers such as Scopus or Web of Science [36]. Unlike Google Scholar, however, Microsoft Academic Graph enables large-scale downloads, and OpenAlex offers API access. The OpenAlex platform aggregates data from Crossref, ORCID, and PubMed, as well as open-access research repositories such as arXiv and Zenodo. OpenAlex also covers papers released as preprints.

9 The relevance score is computed by OpenAlex using machine learning techniques applied to metadata such as abstracts, titles, and citation patterns. Scores above 0.5 indicate concepts that are significant descriptors of the paper’s content rather than peripheral topics.

## Notes

### Competing Interest Statement

The authors have declared no competing interest.

### Summary of Updates

Added new results explaining mechanisms behind our baseline findings.

